# Timing matters: phenological constraints and predation shape Arctic community structure

**DOI:** 10.1101/2025.09.30.679582

**Authors:** Frédéric Dulude-de Broin, Pierre Legagneux, Éliane Duchesne, Maria Belke-Brea, Gilles Gauthier, Dominique Berteaux, Dominique Gravel, Simon Gascoin, Aurore Dupuis, Joël Bêty

## Abstract

Top-down and bottom-up controls of animal populations are key elements of niche and coexistence theories, but there is still little empirical evidence on how these forces determine species distribution and community assemblies. In Arctic ecosystems, spring snowmelt sets the timing and duration of the snow-free period, thereby controlling food availability, while predation often imposes additional constraints on prey species. The relative importance of these abiotic and biotic filters on distribution is also susceptible to vary with body size. Using 10 years of high-resolution data on all major members of an Arctic vertebrate community and their shared predator, we tested how snowmelt timing interacts with predation to shape species occurrence and community structure. Species occurrence declined with later snowmelt dates, with larger-bodied species being particularly constrained by short snow-free periods. Predation further modulated species occurrence, with responses varying according to body mass. Our findings highlight the combined influence of the phenology of food availability and predation as important filters shaping local community structure. Building on species contrasted responses, we propose a conceptual framework for how phenological constraints and predation jointly shape community assembly in highly seasonal environments.

## Introduction

Niche theory is a cornerstone of community ecology and has evolved considerably since original propositions by Grinell (1917) and Elton (1927; see Chase & Liebold (2003), Gravel *et al*. (2019) for reviews). The niche is defined as the set of abiotic and biotic environmental conditions under which a species can persist (Hutchinson 1957), i.e. maintain a positive intrinsic growth rate (Holt 2009). It is a central tenet to both coexistence (Chesson 2000) and species distribution (Godsoe *et al*. 2017) theories. MacArthur’s theory of limiting similarity introduced a bottom-up perspective on niche differentiation, where species coexistence depends on resource partitioning and competitive abilities (MacArthur 1972). In contrast, Holt’s concept of apparent competition introduced a top-down perspective, where species sharing common predators can indirectly affect one another’s abundance and persistence (Holt 1977). These contrasting forces were unified by Chesson & Kuang (2008), who established that species niche overlap and ultimately coexistence should reflect the combination of both bottom-up and top-down interactions. Chesson’s integration of the niche theory has yet to be applied to species distribution theory. Understanding the relative importance of resource availability and predation on species distribution could solve long lasting questions in biogeography and community assembly (Godsoe *et al*. 2017; Kissling *et al*. 2012; Wisz *et al*. 2013).

Energy availability is one of the most fundamental drivers of species community assembly (Lindeman 1942). Plants absorb energy from sun radiation, which is then transferred to other trophic levels by consumptive interactions. Herbivores can establish and persist in a system only if the available energy is sufficient to support their growth and reproduction (Lindeman 1942). Yet, while the total energy available may set an upper limit on species abundance and diversity, the timing of resource availability is also critical. In many biological systems, there is a limited period each year during which environmental conditions are suitable for critical life-history events like growth or reproduction (Hjort 1914; McNamara & Houston 2008; Visser & Gienapp 2019). This phenological window is determined by a combination of favourable abiotic conditions (e.g., adequate pH, favourable temperature, lack of snow) and high resource availability (McNamara & Houston 2008). These windows limit species occurrence based on their physiological tolerances and ability to complete energetically demanding life-history events within the restricted time available (Durant *et al*. 2007; McNamara & Houston 2008). The timing and duration of phenological windows thus filter which species can locally complete their lifecycle and persist (Godsoe *et al*. 2017; Schneider & Frost 1996; Williams *et al*. 2017).

Species vary in how they navigate constraints imposed by phenological windows due to differences in functional traits (Humphries *et al*. 2017). Body mass is a fundamental trait shaping both energetic demands and trophic relationships (Brose 2010; Gravel *et al*. 2013; Riede *et al*. 2011). Because metabolic rate scales with body mass (Brown *et al*. 2004; Kleiber 1932), it is a strong indicator of energy requirements and can serve as a proxy for several other traits (Peters 1986; Sæther 1987). Thus, a species body mass should be intimately linked to the phenological window it needs to persist and reproduce.

Phenological constraints on resource availability can propagate to higher trophic levels, as areas with longer productive periods may support more consumers, in turn altering top-down dynamics (Fretwell & Barach 1977; Hopcraft *et al*. 2012; Oksanen *et al*. 1981). Therefore, longer growing periods tend to support greater prey biomass, which in turn can sustain higher predator abundance (Oksanen *et al*. 1981). Predator density may change nonlinearly and increase rapidly once the growing season is long enough to allow new prey species. This is particularly striking in the case of large colonial prey, such as waterfowl, which can drastically increase predator densities (Dulude-de Broin *et al*. 2023; Giroux *et al*. 2012). The ability of species to withstand predation should also be influenced by their body mass. For instance, the body mass ratio between predators and their prey affects the rate and success of attacks, as larger prey are more difficult to subdue and consume (Brose 2010; Portalier *et al*. 2019). Consequently, phenological constraints not only shape resource availability for primary consumers but may also alter biotic interactions within food webs.

Although the interplay between top-down and bottom-up processes is known to affect population abundance (Hopcraft *et al*. 2010; Hunter & Price 1992), their joint influence on vertebrate community structure and distributions remains much less studied (Godsoe *et al*. 2017; Tylianakis & Morris 2017). In particular, the combined effects of food resource phenology and predation on community structure are rarely integrated (Ponti & Sannolo 2023), likely due to the challenge of obtaining concurrent information on phenological windows, predation risk and multi-species distribution. Arctic terrestrial ecosystems offer a unique opportunity to tackle these questions as seasonal changes impose severe phenological constraints on ecological processes (Humphries *et al*. 2017; Saulnier-Talbot *et al*. 2024). Snow cover creates strong heterogeneity in the length of the growing season (Niittynen & Luoto 2018). In particular, the spring snowmelt unlocks the window of opportunity for plant growth, making it the primary phenological driver of primary production (Walker *et al*. 2001). Snowmelt timing is also intimately linked to the duration of the snow free period, which can constrain consumers’ access to terrestrial food resources (Rixen *et al*. 2022). Here we benefited from a spatial gradient of snowmelt over a 600 km^2^ study area and contrasted predator densities to investigate how the timing and duration of food availability, assessed through spring snowmelt timing, and predator density influence species occurrence and community structure in the High Arctic tundra. We leveraged a 10-years, high-resolution dataset on vertebrate occurrence to: (1) evaluate how spring snowmelt timing influences the occurrence and distribution of 11 species during the breeding period, and (2) assess the response of these species, differing in body mass, to both snowmelt and predation. We hypothesized that snowmelt timing influences local species assemblage by defining the temporal window during which energy is available to complete reproduction. Second, we hypothesized that predator density further modulates species occurrence, and that body mass mediates the joint response to snowmelt and predation (Figure 1). We predicted occurrence would decline with later snowmelt, especially for larger-bodied species with longer, energy-demanding reproductive cycles. However, we expected larger-bodied species to be less sensitive to predation due to greater defence capacity. Together, these hypotheses emphasize the combined influence of abiotic (snowmelt) and biotic (predation) drivers on the structure of vertebrate communities and set the stage for a trait-based examination of species responses across environmental gradients.

**Figure 1.**
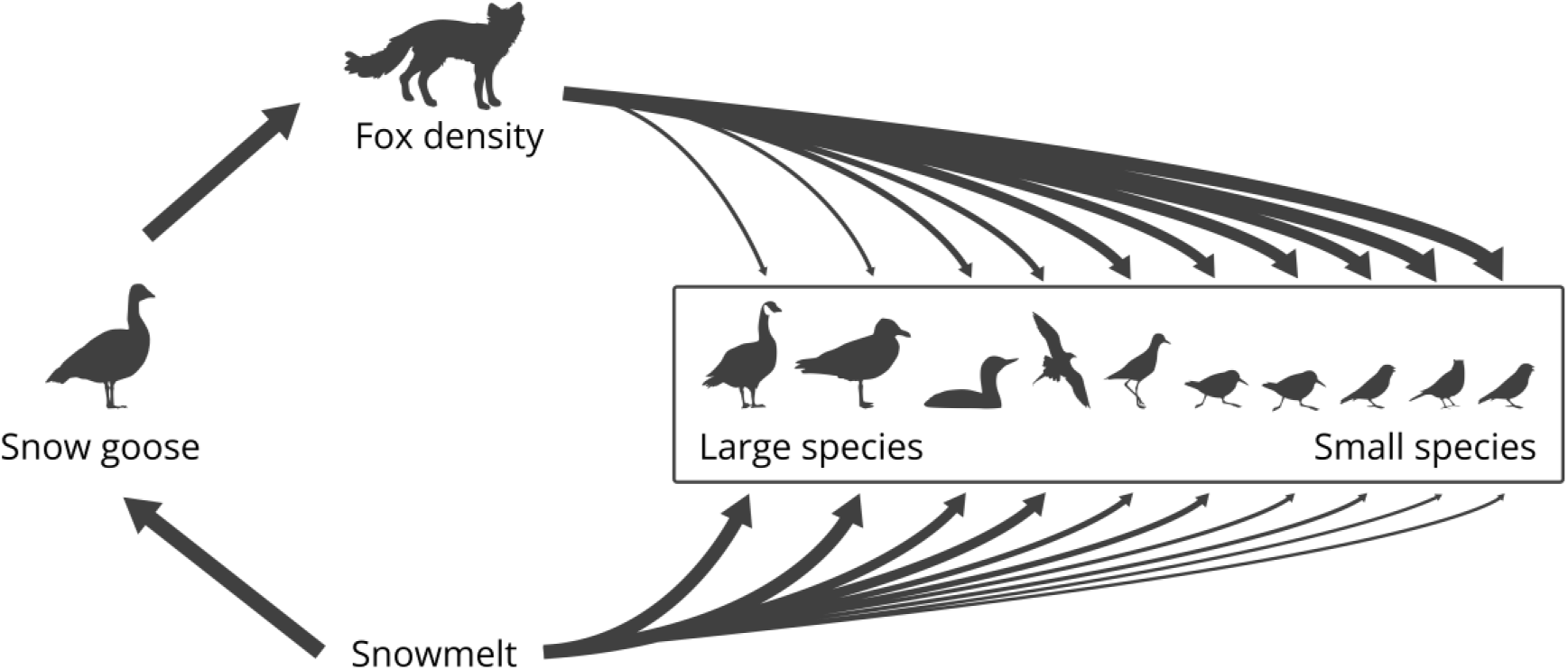
Combined influence of abiotic (snowmelt) and biotic (predation) drivers on Arctic community structure. We hypothesize that snowmelt timing sets the temporal window for reproduction, influencing species occurrence, while predator density further modulates these patterns. Body mass is expected to mediate species responses, with larger-bodied species more sensitive (wider arrow) to snowmelt timing but less affected (narrower arrow) by predation. Note that the abundance of all prey species can influence predator density, but the presence of large colonial prey, such as snow geese, can have a dominant effect.

## Materials and Methods

### Study system

Field work was conducted on Bylot Island (72.93°N, 79.76°W), in Sirmilik National Park, Nunavut, Canada (Figure 2). The ∼600 km² area (Gauthier *et al*. 2011) is mostly covered by mesic tundra, with wet meadows commonly found in valley bottoms (Gauthier *et al*. 2011). Lakes and ponds are scattered throughout the landscape. Elevation increases from the coast and valley bottoms to an inland plateau, up to 600m asl. Mesic tundra and wet meadows are well vegetated and rocky substrates associated with xeric tundra become more common with altitude. Nonetheless, all major habitat types are broadly distributed throughout the study area. A large snow goose (*Anser caerulescens*) colony occupies a ∼70 km^2^ area where around 5,000 to 20,000 pairs nest each year (Gauthier *et al*. 2011).

**Figure 2.**
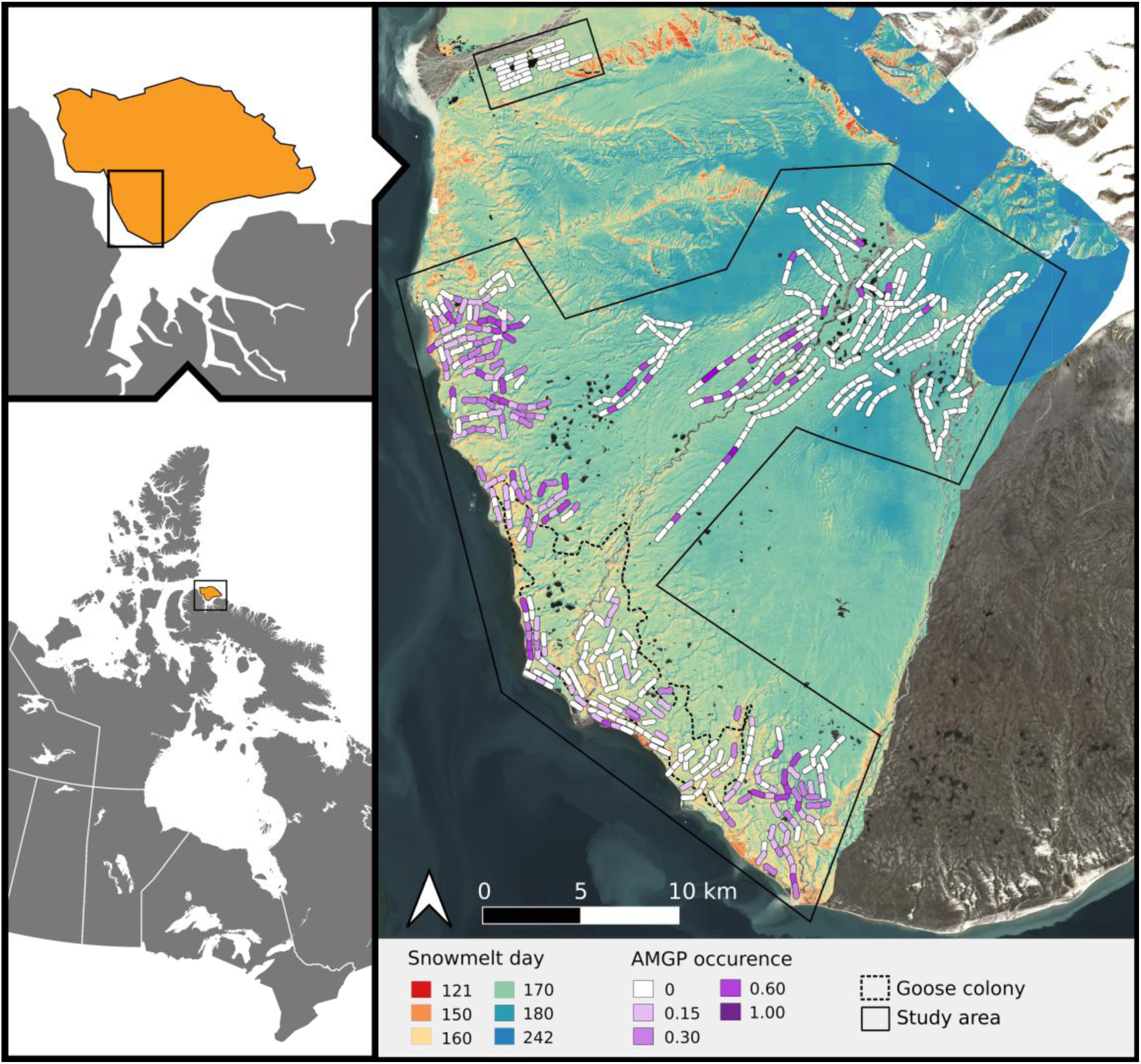
Location of the Bylot Island study area (left panels). Right panel shows the study zone (inside solid black lines), the location of sampling transects (segmented lines), the greater snow goose colony boundary (black dashed line), and average snowmelt timing. The occurrence of studied species was recorded within 150 m on each side of 500 m-long transects or around lakes. For illustrative purposes, segment colours indicate the relative occurrence (i.e., the proportion of surveys with a sighting) of the American golden plover (AMGP), one of 11 surveyed species. Snowmelt day refers to the day of the year (counting from January 1) and was derived from combined Sentinel-2 and MODIS imagery. Areas where snow did not melt were assigned a value of 242.

Arctic fox is the main terrestrial predator and feeds primarily on lemmings (brown lemming, *Lemmus trimucronatus*, and collared lemming, *Dicrostonyx groenlandicus*) and snow goose eggs, but also opportunistically consume the eggs of other tundra nesting birds (Gauthier *et al*. 2011). We assessed the occurrence of 11 bird species, which share the arctic fox as primary predator: greater snow goose, cackling goose (*Branta hutchinsii*), glaucous gull (*Larus hyperboreus*), red-throated loon (*Gavia stellata*), long-tailed jaeger (*Stercorarius longicaudus*), American golden plover (*Pluvialis dominica*), Baird’s sandpiper (*Calidris bairdii*), white-rumped sandpiper (*Calidris fuscicollis*), Lapland longspur (*Calcarius lapponicus*), horned lark (*Eremophila alpestris*), and snow bunting (*Plectrophenax nivalis*). These species vary in body size, nesting habitat, diet, and vulnerability to fox predation (See Appendix S1 for detailed descriptions of these species). Ground-nesting shorebirds and passerines are highly vulnerable to fox predation (McKinnon & Bêty 2009), whereas larger and water-associated birds have partial defences and may benefit from nesting microhabitats less accessible to foxes (Corbeil-Robitaille *et al*. 2024).

### Nesting bird occurrence

We conducted surveys along transects to assess the occurrence of species commonly found nesting in mesic and xeric habitats, and we surveyed lake shores to assess the occurrence of species nesting near lakes and ponds.

### Transects

Observers walked at constant speed along 500 m transects and recorded all bird sightings, along with reproductive status (see Appendix S1 for breeding criteria), within 150 m on either side (distances estimated by observers trained with range finders). For each species, detection probability was assumed constant across observers and habitats, given the use of standardized protocols, trained observers, and the open tundra landscape. Most species were highly detectable often responding to observers’ presence, and the open tundra provided excellent visibility, minimizing potential influence of habitat structure on detection rates. Surveys along permanent transects were carried out annually between 2010 and 2024, except in 2020 and 2021, with an average of 243 transects per year (range: 146-295). Additionally, from 2022 to 2024, we added ∼300 transects in areas with later snowmelt. For all years but 2024, surveys took place during bird incubation period from 20 June to 14 July (see Appendix S1). We alternated surveys between early and late snowmelt areas, as well as between areas with high and low fox densities (see below), to minimize bias from potential seasonal decline in nest detection due to nest failure (Lamarre *et al*. 2017). .

For analyses, we used the occurrences of breeding individuals along transects for plovers, jaegers, and passerines, and both breeding and unknown-status individuals for other species, excluding non-breeders. Only high lemming years were included for jaegers as they do not nest at low lemming density (Seyer *et al*. 2023). Snow bunting analyses focused on data from 2022 and 2023, as the addition of transects in late snowmelt areas provided most observations for that species.

### Lake surveys

Lakes across the study area were surveyed for nesting cackling goose, glaucous gull, and red-throated loons from 2022 to 2024 (n=288 lakes in 2022, 399 in 2023, and 117 in 2024). Observers walked the perimeter of each lake, carefully searching for nests along the shoreline and on islets, to record presence or absence of a nest for each species.

### Nesting phenology

For snow geese, American golden plovers and Lapland longspurs, intensive nest searches independent of transect surveys were conducted, providing large samples of laying dates (snow goose: n=8496 over 27 years, American golden plover: n=566 over 15 years, Lapland longspur n=766 over 25 years). For those species, we present the distribution of laying dates alongside the occurrence-snow relationships.

### Snowmelt phenology

To assess the influence of snowmelt patterns on spatiotemporal variation in species occurrence, we mapped the date of spring snow disappearance (hereafter snowmelt date) by calculating, for each year from 2010 to 2024, the median snowmelt date of all pixels within 150 m of surveyed transects and lakes. We focused on snowmelt date rather than snow cover duration because it is the dominant phenological driver of primary production in the Arctic (Walker *et al*. 2001). However, both were highly related (Appendix S1, Figure S10).

Snowmelt dates can be derived from Sentinel-2 imagery from 2017 onward (spatial resolution: 20 m, revisit frequency: 1-2 d on Bylot, Drusch *et al*. 2012) and from MODIS data between 2000 and 2024 (resolution: 500 m, revisit frequency: 1 d on Bylot, Hall & Riggs 2021). To benefit from both the high spatial resolution of Sentinel-2 products and longer temporal coverage of MODIS, we produced enhanced snowmelt maps by combining information from both satellites. Methodological details and validation are available in Appendix S1.

### Predator density

Arctic fox density varies within the study area, largely due to the presence of the snow goose colony. Home ranges of foxes with dens located within the goose colony are half the size of those outside the colony, resulting in higher local densities (Dulude-de Broin *et al*. 2023). To quantify variations in fox local density, we used spatial variations in home range size compiled from fox tracking data (Berteaux 2020, 2021). These tracking data allowed us to delineate 248 individual fox home ranges over 16 years, which collectively overlapped with 56% of transects and 75% of lakes surveyed (see Appendix S1 for analysis of tracking data).

For transects and lakes with spatially overlapping home ranges (n_transect_=372, n_lake_=332), we calculated the average size of fox home ranges, regardless of the year. Estimates were based on a mean of 16 overlapping home ranges per transect (range: 3-39) and 15 per lakes (range: 1-35). Home range analysis allowed the classification of transects and lakes in two categories, large and small home ranges, largely related to fox access to the goose colony (see Appendix S1). Transects and lakes that did not overlap with documented home ranges (n_transect_=288, n_lake_=112) were assigned to the large home range size category because they were at least 10 km from the goose colony. To facilitate interpretation, we refer to high and low predator density throughout the manuscript.

### Lemming density

We used annual lemming abundance as a covariate in our analysis. Lemmings were live-trapped in two 11-ha permanent grids, one in wet and one in mesic habitat, to obtain spatially explicit mark–recapture estimates of density (methods described in Fauteux *et al*. (2015)). Trapping sessions were conducted in June, July and August and we used the average density for all three sessions and both lemming species to obtain annual lemming density. We then classified each transect and lake survey into one of two classes of lemming density, that is low (<18 lemming/km^2^) or high (>174 lemmings/km^2^). No intermediate lemming density was observed during the study period. Moreover, these two lemming density categories closely reflect fox breeding propensity throughout the study area (Giroux *et al*. 2012; Juhasz *et al*. 2020).

### Statistical analyses

For each bird species, we identified the latest snowmelt date compatible with reproduction (hereafter snowmelt threshold), defined as the latest snowmelt date across all transects or lakes with a nesting individual. We quantified uncertainty using bootstrap resampling (see details in Appendix S1). We modelled the influence of snowmelt date and predator density on the probability of occurrence of a species on individual transects and lakes using generalized linear mixed models with a binomial distribution and a logit link. Each species was analysed separately, and we used model selection to evaluate the contribution of each variable.

Our analysis followed a two-step approach. First, because we expected non-linear relationships between snowmelt dates and the probability of occurrence, we fitted the full model (see below) using natural cubic splines with degrees of freedom ranging from 1 to 4. The most parsimonious model was selected based on AICc. Second, we compared a set of models with different combinations of snowmelt dates (fitted with the selected spline), predator and lemming density (both, high vs. low), and the interaction between predator and lemming density. All models included year as a random intercept. Models based on transect also included transect ID as a random intercept to account for repeated surveys over the years. Lake ID was not included because nearly half the lakes (42%) were surveyed only once. We considered models with ΔAICc< 2 to be equally supported (Burnham & Anderson 2002). To illustrate the effect, or lack of effect, of the main variable of interests (snowmelt date and fox density), we present predictions from the most parsimonious model that includes both variables, rather than from the best-fitting model. All predictions are shown for high lemming density.

We evaluated how response to snowmelt timing varies with body mass by comparing snowmelt thresholds across species. We similarly assessed how responses to predation varies with body mass by comparing the effect sizes of predator density extracted from the full models, which accounted for snowmelt timing, across species.

## Results

We monitored 637 unique transects and 444 lakes resulting in 4097 transect-years and 804 lake-years (sightings counts by species are provided in Appendix S1). Median snowmelt dates ranged from day 140 (May 20) to day 203 (July 22) for transects and from day 155 (June 4) to day 194 (July 13) for lakes. On average, snowmelt timing varied by 18 days across years for the same transect and by 9 days for the same lake. Within a given year, snowmelt timing varied on average by 24 days among transects and 29 days among lakes.

Transects and lakes were well distributed in terms of predator density with 30% of transects and 43% of lakes monitored in areas with high fox densities, and 48% and 15% surveyed during years of high lemming density, respectively. The occurrence of foraging foxes on transect was on average 4.2 times higher in areas of high fox densities compared to areas with low fox densities (Appendix S1, Figure S1).

Occurrence probability of nesting species was generally high in early snowmelt areas and declined with later snowmelt dates until reaching a threshold beyond which a species was no longer observed (Figure 3-4, see Appendix S1, Table S1 for threshold values). Waterfowls had the earliest snowmelt thresholds and did not nest in areas still covered by snow in late June, a pattern also observed in gulls and jaegers. Red-throated loons and American golden plovers nested in regions with later snowmelt, while smaller species nested across the study area, with reproductive occurrences declining only under very late snowmelt (Figures 3-4). Notably, passerine thresholds were near the latest snowmelt dates sampled. For species with intensive nest search data, snowmelt thresholds closely matched the latest observed lay dates (Figure 3). As expected, larger-bodied species were most affected by a late snowmelt and had the earliest thresholds (Figure 5A).

**Figure 3.**
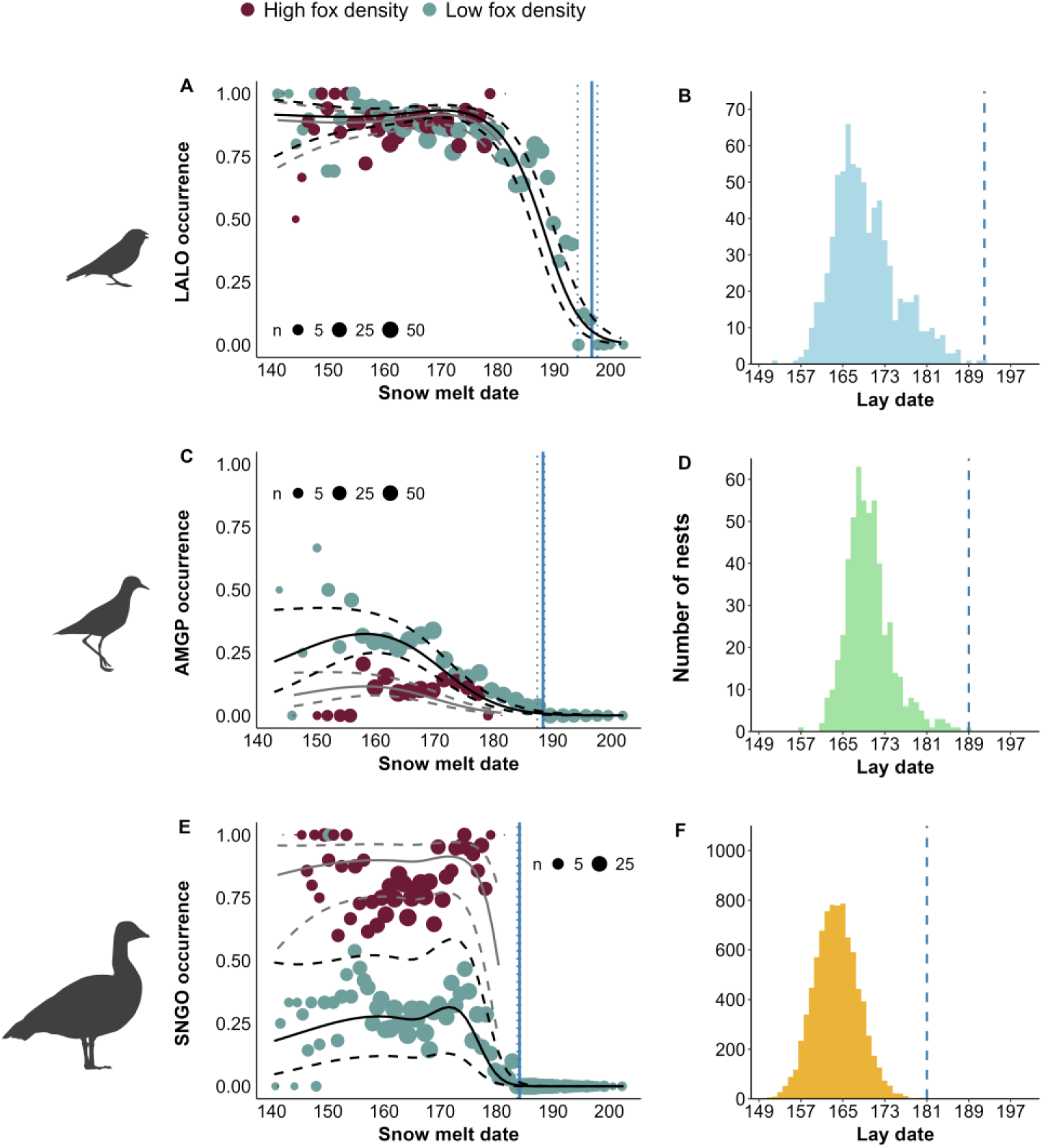
Influence of snowmelt date on occurrence probability of breeding (A) Lapland longspur (LALO), (B) American golden plover (AMGP), and (C) greater snow goose (SNGO) on Bylot Island, Nunavut (2010–2024). The full and dotted lines show model predictions along with their 95% confidence intervals for transects located in low (black) or high (grey) fox density areas. The dots are raw species detection averaged over equally sized intervals for transects located in low (green) and high (dark red) fox density areas. Circle size is proportional to log(n). Snowmelt date refers to the median snowmelt date within 150 m on each side of the 500 m transects. The vertical solid and dashed lines are the snowmelt threshold with the 95% CI. For each species, the distribution of nest laying dates is shown to illustrate correspondence with the snowmelt threshold (vertical dashed lines on these graphs; panels B, D, F).

**Figure 4.**
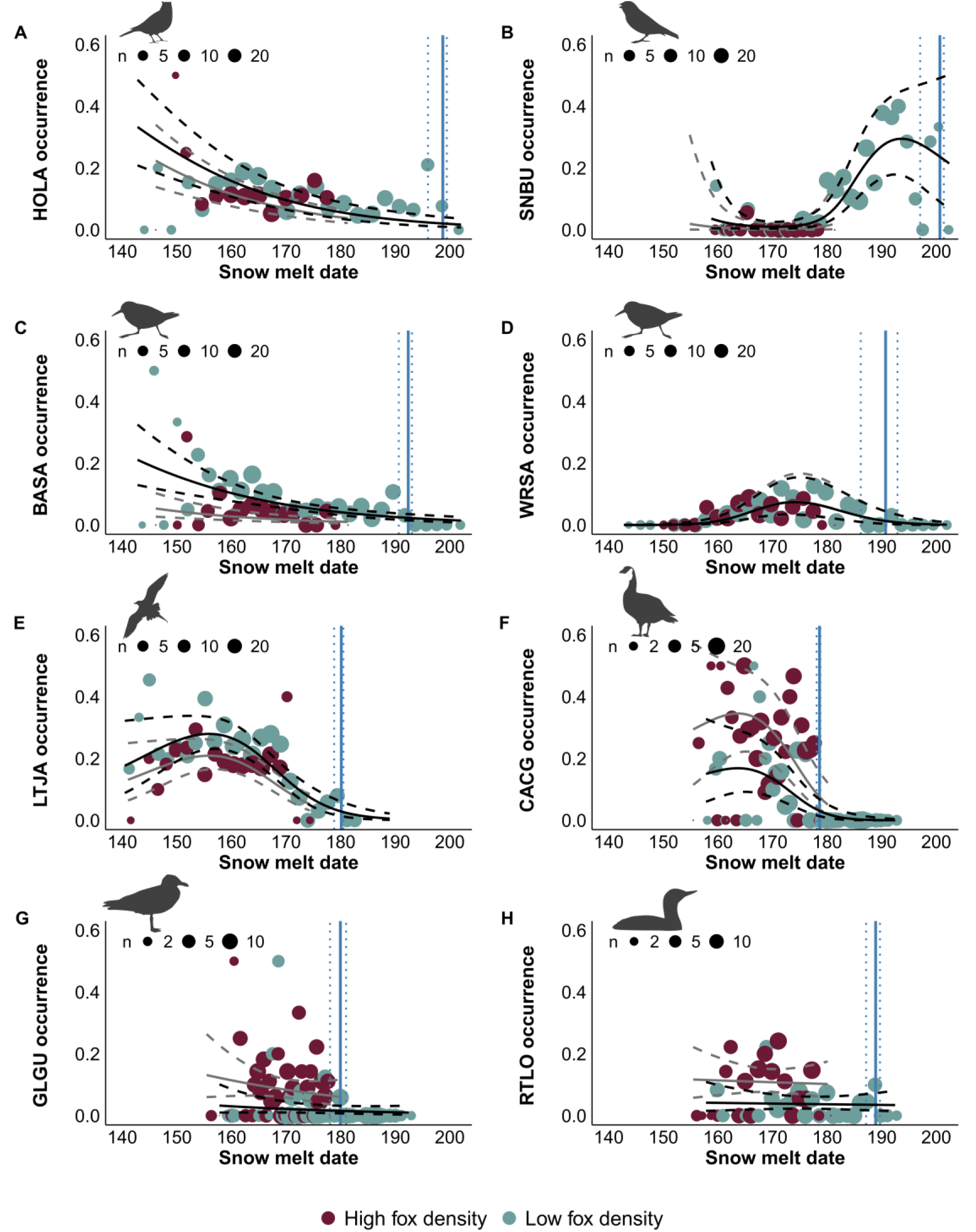
Influence of snowmelt date on occurrence probability of breeding (A) horned lark (HOLA), (B) snow bunting (SNBU), (C) Baird’s sandpiper (BASA), (D) white-rumped sandpiper (WRSA), (E) long-tailed jaeger (LTJA), (F) cackling goose (CACG), (G) glaucous gull (GLGU), and (H) red-throated loon (RTLO) on Bylot Island, Nunavut (2010–2024). The full and dotted lines show model predictions along with their 95% confidence intervals for transects or lakes located in low (black) or high (grey) fox density areas. The dots are raw species detection averaged over equally sized intervals for transects located in low (green) and high (dark red) fox density areas. Circle size is proportional to log(n). Snowmelt date refers to the median snowmelt date within 150 m on each side of the 500 m transects (species A-E) or around surveyed lakes (species F-H). The vertical solid and dashed lines are the snowmelt threshold with the 95% CI.

**Figure 5.**
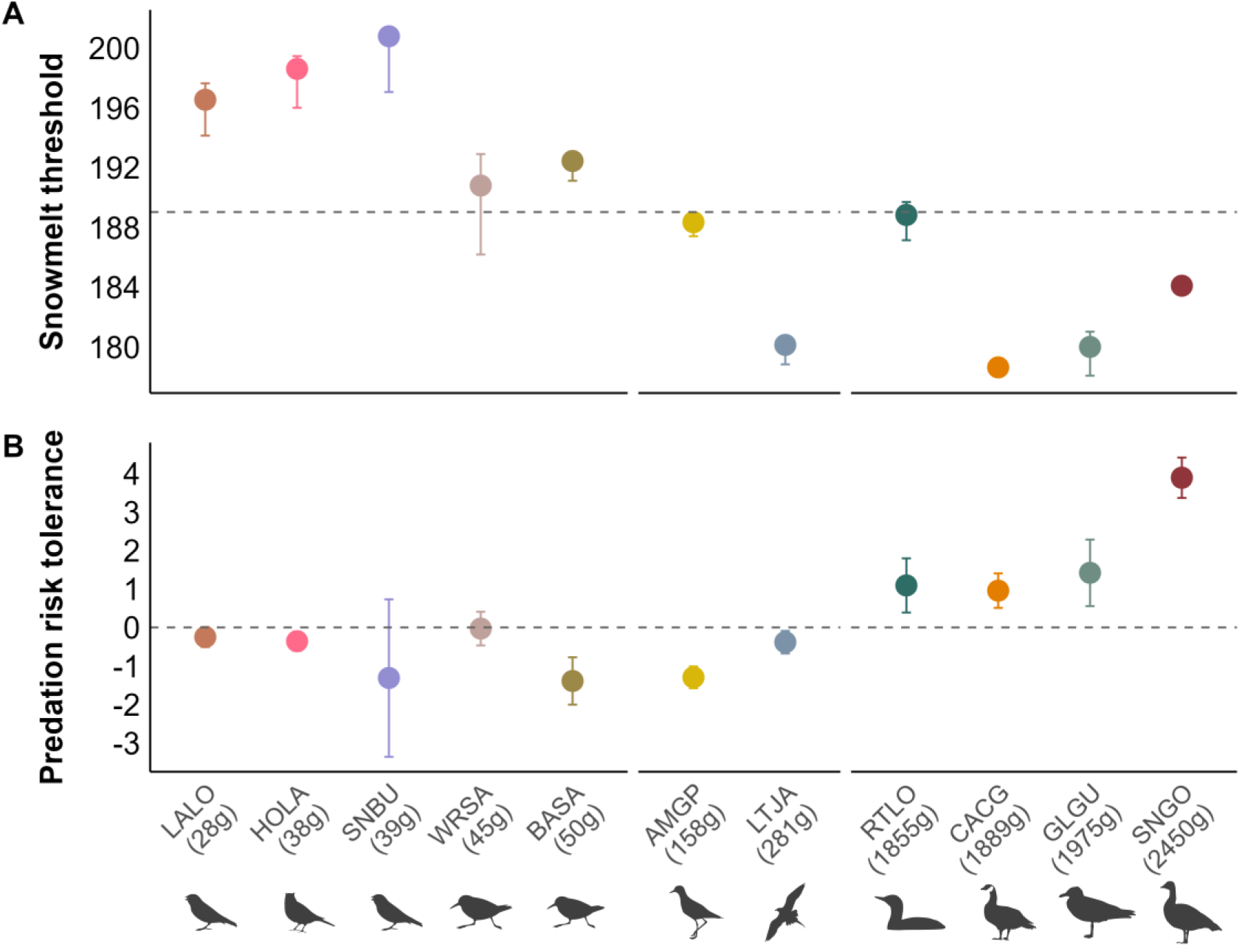
Snowmelt threshold dates for reproduction and predation risk tolerance of species ordered by body mass. (A) Snowmelt threshold corresponds to the last date a breeding individual was observed. Dots show the threshold for each species with 95% confidence intervals from non-parametric bootstrap. (B) Predation risk tolerance is the effect of fox density on species occurrence probability (values in Appendix S1, Table S3), with dots showing effect sizes and 95% confidence intervals. Negative predation risk tolerance indicates lower probability of occurrence in high fox density area.

Snowmelt date was an important predictor of occurrence in all species except glaucous gulls and red-throated loons (Figure 3-4) for which snow was not included in the most parsimonious model (Appendix S1, Table S2). Occurrence probability of Lapland longspurs, American golden plovers, long-tailed jaegers, cackling geese and snow geese was high in early snowmelt areas and declined steeply in late snowmelt areas, while the decline was more gradual for horned larks and Baird’s sandpipers (Figures 3-4). White-rumped sandpipers showed a humped shaped pattern, with low occurrence at early and late snowmelt sites. Notably, snow buntings occurred mainly in sites where snowmelt happened after day 180, before declining again.

Predator density was an important predictor of occurrence probability as fox density was included in the most parsimonious model for most species (Appendix S1, Tables S2). However, associations were either positive or negative depending on species (Figures 3-4). As expected, snow geese occurrence was positively associated with areas of high fox density (Odds ratio (OR)=140.85 [72.74, 272.73], 95% CI). The same trend was found in cackling geese (OR=2.47 [1.57, 3.87]), glaucous gulls (OR=3.99 [1.67, 9.53]), and red-throated loons (OR=3.62 [1.69, 7.74]). In contrast, Horned larks (OR=0.70 [0.55, 0.90]), Baird’s sandpipers (OR=0.25 [0.13, 0.46]), American golden plovers (OR=0.28 [0.21–0.37]), long-tailed jaegers (OR=0.68 [0.51, 0.91]) and, marginally, Lapland longspurs (OR=0.78 [0.60, 1.01]) were less likely to occur in areas of high fox density (22% to 74% reduction in probability of occurrence). Fox density did not affect the occurrence of white-rumped sandpipers and snow buntings (Tables S2 and S3). Lemming density had a lesser effect on species occurrence than fox density as it was in the most parsimonious model only for snow goose and Baird’s sandpiper (Appendix S1, Table S2).

Large-bodied species (≥1855 g) were generally less sensitive to predation than smaller ones (Figure 5B), showing positive associations with areas of high fox densities. Interestingly, although smaller species were generally negatively affected by fox densities, the strongest effects occurred among species of intermediate body size (50-150 g; Figure 5B).

## Discussion

Benefiting from a 10-year dataset on 11 bird species and their predator across 600 km^2^ of Arctic tundra, we found that species occurrence declined with later snowmelt until a threshold beyond which breeding was no longer observed, with the strongest effects observed in larger-bodied species. The occurrence of breeding species was further shaped by predation, with responses varying according to body mass. Our findings highlight the combined influence of the phenological timing of resource availability and predation as fundamental ecological filters structuring vertebrate communities.

### Snowmelt timing

Our results support the hypothesis that local assemblages of breeding species depend on the phenological windows set by snowmelt timing. The ecological effects of snowmelt are well documented in boreal and Arctic systems (Slatyer *et al*. 2022).

Earlier snowmelt has been linked to earlier reproduction (Dickey *et al*. 2008; Liebezeit *et al*. 2014) and higher breeding success (Anderson *et al*. 2015; Keyser *et al*. 2022; Post & Forchhammer 2008) potentially driving species distribution (Keyser *et al*. 2022). In extreme cases, abrupt changes in snowmelt timing can trigger ecosystem-wide reproductive failure (Schmidt *et al*. 2019). Together with these studies, our results underscore snowmelt as a key driver of tundra community structure.

Food availability duration constrained by snow cover was likely more limiting for nesting species than total available biomass from primary production. At early snowmelt sites, where herbivory pressure is the highest, only a small fraction (<10%) of primary production is consumed annually (Legagneux *et al*. 2012). This suggests that other processes than total food availability are limiting. The issue is likely not how much, but rather how long these resources are available. Similar mechanisms could shape community structure in other highly seasonal systems where reproduction is tightly linked to food temporal availability. One such example is songbird communities relying on synchronized insect emergence (e.g. Visser & Both, 2005).

Species had a clear snowmelt threshold for reproduction as observed in other systems (Smith *et al*. 2010), and large-bodied species were generally more sensitive to late snowmelts. Our results support the hypothesis that large species are constrained by the duration and timing of resources availability, because their higher energetic demands and longer developmental periods prolong their vital cycle. However, other traits such as diet and energy storage capacity may also shape species-specific responses. For example, red-throated loons had thresholds that did not align well with their mass. Species phenological tolerance should be closely tied to their food resources (Altermatt 2010; Diamond *et al*. 2011). Therefore, species such as loons (Davis 1972) and gulls (Gauthier *et al*. 2015) that can forage at sea might be less affected by terrestrial snowmelt than herbivores or terrestrial insectivores. Energy stores or the ability to forage under the snow could also extend species’ phenological window by providing access to resources during less productive periods (Kerby & Post 2013; Williams *et al*. 2017). For instance, geese would likely need longer pre-breeding foraging period without body stores. To minimize potential biases associated with imperfect detection and nest failure, we focused on presence/absence data over broad areas and alternated surveys across snowmelt and predator gradients. However, differences in detectability across species are likely and occurrence probabilities should not be directly compared. Importantly, detectability differences are unlikely to have biased observed snowmelt thresholds, as the larger, more detectable species showed earlier thresholds than the smaller, more cryptic ones. Imperfect detection would advance threshold estimates, rather than delay them, making our observed patterns conservative.

In addition to the direct impact of snowmelt timing on resource availability, other processes could contribute to patterns of species occurrence. Prolonged snow cover duration can impact habitat types by altering soil dynamics and plant communities (Niittynen *et al*. 2018, 2020). Although surveys were designed to minimize habitat differences, fine-scale vegetation variation may still locally impact occurrence. For example, white-rumped sandpipers were rare in the earliest snowmelt areas, potentially reflecting a preference for upland sedge meadows (Smith *et al*. 2007). Similarly, snow buntings occurred only in late-melt areas, consistent with their reliance on rock cavities (Montgomerie & Lyon 2020) mainly found in areas where prolonged snow cover exposes rocky ledges by limiting vegetation growth. These associations could suggest an indirect link between snow-free period length and nesting habitat availability or reflect underlying factors that influence both snowmelt and habitat. Other processes such as colonial nesting or site fidelity may also decouple occurrence from annual snowmelt dynamics. Comparing the explanatory power of annual and long-term averaged snowmelt dates could help clarify the timescale over which vertebrate communities respond to these snowmelt gradients. Still, the inverse snowmelt-body mass relationship we observed points to effects mediated by primary production and energy availability, rather than habitat composition. Overall, our results suggest that snowmelt timing play a central role in shaping community structure.

#### Predation

Predation further shaped the occurrence of most nesting birds although its influence varied depending on species. Previous studies in this (Beardsell et al., 2023; Clermont et al., 2021; Duchesne et al., 2021; Dulude-de Broin et al., 2023; Lamarre et al., 2017) and other Arctic tundra ecosystems (Etchart *et al*. 2025; Flemming *et al*. 2019) have shown that predation is a strong driver of nesting success that could shape breeding bird distribution. Within our study area, early snowmelt sites host abundant food resources for predators provided in large part by the presence of a goose colony, which increases fox density and thus predation risk for prey (Dulude-de Broin *et al*. 2023). American golden plovers and Baird’s sandpipers were the species most affected by high fox density, likely because they are large enough to facilitate detection, but cannot efficiently defend their nest (Beardsell et al. 2021). The negative effect of predation was also clear for long-tailed jaegers, which are highly detectable and have limited defences against foxes. Smaller passerines cannot defend their nests either, but they may be harder to detect (Beardsell *et al*. 2021) and their shorter breeding cycles (20-25 days) than shorebirds (36-55 days) reduces their exposure to predation.

In contrast with previous negative associations, the occurrence of the largest species (gull, loon, and geese) was positively associated to areas of high predator density. For colonially nesting snow geese, the aggregation, predictability, and storage potential of eggs attract foxes to the colony and shift cost-benefit trade-offs towards the formation of smaller home ranges, resulting in higher fox densities (Dulude-de Broin *et al*. 2023). Snow goose eggs also provide additional food resources for gulls (Gauthier *et al*. 2015), which may promote nest formation in areas of high snow goose and, indirectly, high fox densities. In addition, attacking gull or goose nests may be risky for foxes and they appear to decrease risk-taking behaviours when their energy acquisition rate is high (Beardsell *et al*. 2025). The positive associations suggests that these species can tolerate high fox densities likely because they can effectively defend their nest or use partial refuges (e.g. islets) to reduce predation risk (Beardsell *et al*. 2025; Beaudoin 2025; Corbeil-Robitaille *et al*. 2024). These potential mechanisms illustrate how abiotic constraints (snowmelt timing) can rapidly propagate and create complexity, even within simple food webs.

We found limited effect of lemming availability on species occurrence. While high lemming densities are known to reduce nest predation (Beardsell *et al*. 2022; Bety *et al*. 2002; McKinnon *et al*. 2014), the reduction it induces (0.5-1 fold; Beardsell et al., 2022; McKinnon et al., 2014) is small compared to the sharp increase driven by fox numerical response near the goose colony (up to six-fold; Dulude-de Broin et al., 2023). Our monitoring was based on species presence-absence along transects or lake shores rather than abundance, which may have limited our ability to detect such smaller predator release at high lemming density.

In our study system, the snow goose colony strongly amplified predation pressure, which helped assess how variation in predator density influences species occurrence across the landscape. Early snowmelt was necessary for the establishment of such a colony, but not all early snowmelt areas would experience the same level of predation in the absence of snow geese. Still, the relative occurrence of other nesting species (e.g. cackling goose, gull, loon, plover, jaeger) indicates that combined prey biomass could be higher in early snowmelt areas even in the absence of a colony. Early snowmelt could also promote predator access to prey sheltered by the snow pack, such as lemmings (Bilodeau *et al*. 2013). Such increases in prey availability driven by snowmelt timing would thus positively influence predator density.

#### Proposed framework

We propose that, once biogeographic constraints are considered, the phenological window of resource availability acts as a secondary ecological filter shaping community structure, setting the stage for a third filter driven by predation (Figure 6). Together, these interacting filters shape the occurrence and relative abundance of nesting species according to their traits, particularly body mass. In late snowmelt areas, the short breeding window excludes larger species that require more time and energy to complete their breeding cycle. In contrast, early snowmelt areas may support more species, including large-bodied and colonial herbivores, which contribute to a higher overall prey biomass. Elevated prey biomass can in turn support greater densities of predators, such as arctic foxes, resulting in stronger predation pressure, which further shapes community composition. However, the effect of predation varies among species due to their traits. Large species that can defend their nest, and small cryptic species that can reduce detection should be less affected by predation. In contrast, those of intermediate body size appear most vulnerable, being both unable to defend their nest and highly detectable. We therefore expect early snowmelt areas to be characterised by larger species, higher overall prey biomass, longer food chain and less vulnerable species, while late snowmelt areas host species more susceptible to predation but less constrained by food availability.

**Figure 6.**
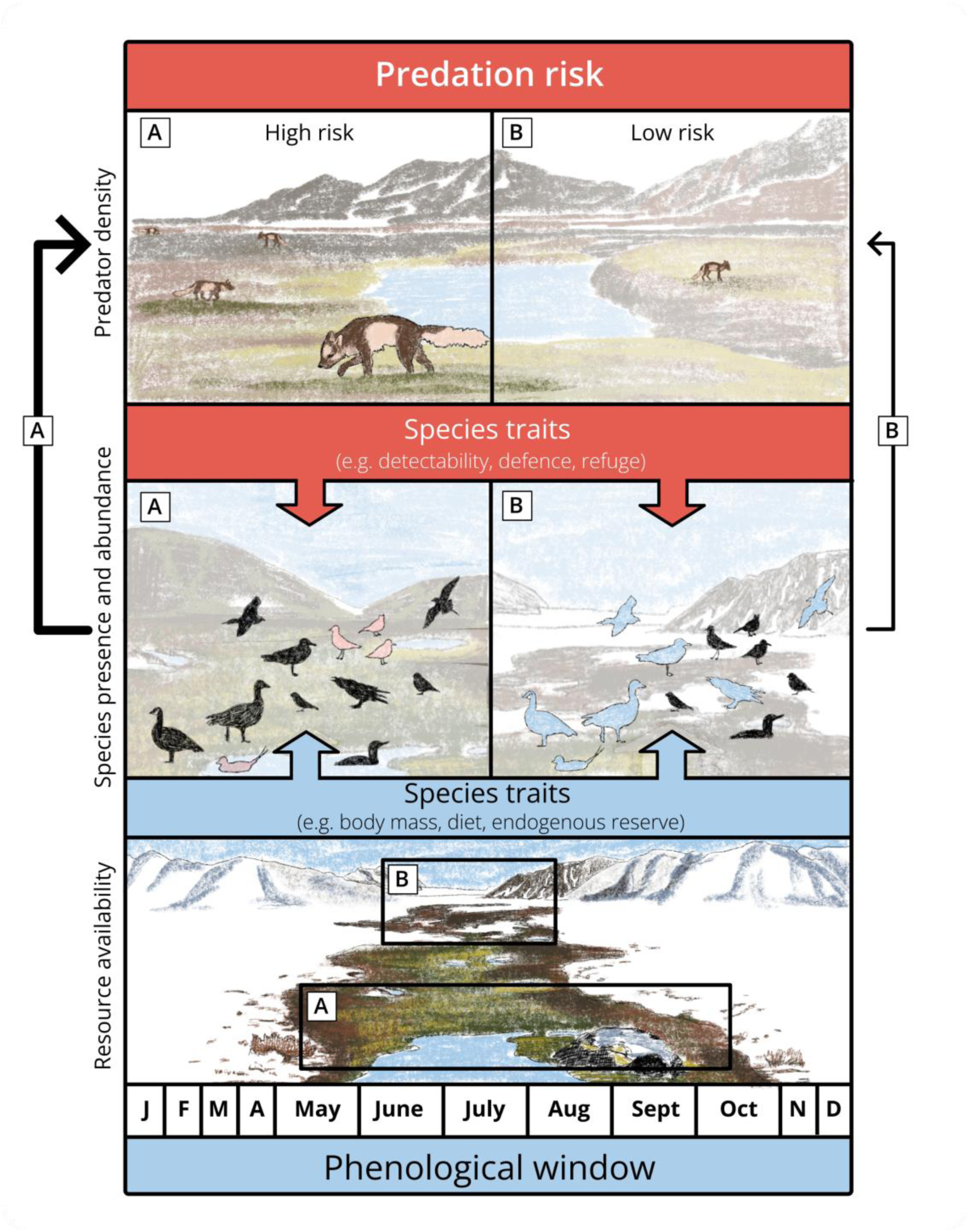
Conceptual model illustrating how phenological constraints and predation shape community composition. Early snowmelt sites (A) allow a longer period of resource availability that can support a broad range of breeding species including those with large body size. This results in higher overall prey biomass than later snowmelt areas, which in turn sustains more predators. Predation then acts as an additional ecological filter, excluding the most vulnerable species (in red) or reducing their abundance. In contrast, sites with shorter phenological windows (B) exclude larger species (in blue) but still offer sufficient resources for smaller species. These smaller species, including those that are more vulnerable to predation, may persist in late snowmelt areas because the absence of larger species reduces predator densities.

## Conclusion

Our study demonstrates that both the timing of snowmelt, a variable strongly associated with the duration of primary production, and predation act as key ecological filters shaping avian communities in the Arctic tundra. We show that the timing and duration of resource availability can impose sharp constraints on breeding opportunities especially for larger species, while intermediate body size species could be most vulnerable to predation. These ideas may extend more broadly to other seasonal systems, providing a framework for understanding how phenological constraints on primary production can propagate through food webs via biotic interactions and shape ecological communities.

## Acknowledgements

We thank Andreanne Beardsell, Mathilde Poirier, Frédéric Letourneux, Alexandra Langwieder, Matthieu Weiss-Blais, and Jeanne Clermont for insightful discussions and comments on the study. We are grateful to Marie-Christine Cadieux and Marie-Jeanne Rioux for their essential support in coordinating fieldwork campaigns and data management. We thank Thierry Grandmont, Éléonore Douville, Laurence Gagnon, Madeleine-Zoé Corbeille-Robitaille, Joassie Ootovak, Laurianne Dumont, Marylou Beaudoin, Louis-Pierre Ouellet, Mathieu Archambault, and the many people who collected data on Bylot Island for this project. We are grateful to the Sirmilik National Park of Canada and the community of Mittimatalik for their support. This work was financially supported by the Canada Foundation for Innovation, the Canada Research Chairs Program, the Kenneth M Molson Foundation, the Natural Sciences and Engineering Research Council of Canada (NSERC), the Canada First Sentinel North research program, the Network of Centres of Excellence of Canada ArcticNet, the Northern Scientific Training Program (Polar Knowledge Canada), and the Polar Continental Shelf Program (Natural Resources Canada). F. Dulude-de Broin received scholarships from NSERC, the Fonds de Recherche du Québec, and Sentinel North program.

## Conflict of Interest

None.

## Author Contributions

Conceptualization: FDB, PL, JB, Formal analysis: FDB, Data curation: FDB, ED, MBB, Visualization: FDB, ED, JB, Writing – original draft: FDB, Writing – review & editing: FDB, PL, ED, MBB, GG, DB, DG, SG, AD, JB, Funding acquisition: PL, GG, DB, JB, Supervision: PL, JB

## Data availability

Data and code will be published in a public repository upon manuscript acceptance.

## Supporting Information

### 1. Additional methods

#### 1.1 Species habitat, diet and vulnerability to predation

Shorebirds and passerines (i.e. Lapland longspur (28g), snow bunting (39g), White-rumped sandpiper (45g), Baird’s sandpiper (50g), American golden plover (158g)) nest in mesic to xeric tundra and feed primarily on arthropods along with seeds and berries (Baker 1977; Leung *et al*. 2018; Montgomerie & Lyon 2020). Long-tailed jaegers (281g) also nest in these habitats and, during the breeding period, mainly feed on lemmings but also on passerines and shorebirds chicks, arthropods, and berries (Andersson 1976; Seyer *et al*. 2020). In contrast, water-associated species such as the cackling goose (1889g), glaucous gull (1975g), and red-throated loon (1855g) nest around lakes and wetlands. Cackling geese feed mainly on vegetation, gulls are opportunistic predators and scavengers eating fish, invertebrates and eggs of other nesting birds (Gauthier *et al*. 2015), and loons primarily consume fish and invertebrates (Davis 1972). Snow geese (2450 g) are colonial nesters that occupy both mesic tundra and wetlands, often concentrating in large colonies but also nesting individually away from the colony center. Their diet consists mainly of vegetation such as grasses, sedges, and forbs. Artic fox (2500g) predation is the main terrestrial predator of all all these species (Gauthier *et al*. 2011; McKinnon & Bêty 2009). Ground-nesting shorebirds and passerines are highly vulnerable to arctic fox predation, whereas larger and water-associated birds have partial defences and may benefit from nesting habitat that reduces fox foraging effort or success (Corbeil-Robitaille *et al*. 2024).

### 1.2 Delayed field work in 2024

In 2024, a helicopter crash delayed all field work activities, and sampling exceptionally occurred from July 21 to August 1, a period that overlaps brood rearing for many species. Although the scale of the analysis exceeds the typical movement range of most focal species during brood rearing, snow geese often engage in long-distance movements during this period (Mainguy *et al*. 2006) and were excluded in 2024. Data for snow bunting and white-rumped sandpiper from 2024 were also excluded because both species become much more detectable during brood rearing due to the presence of large fledglings, and because of the conspicuous distraction displays of brood-rearing white-rumped sandpipers. To further reduce potential bias on that year, we deployed field teams simultaneously across the study areas and completed transects within a condensed 11-day period. We also confirmed that the direction and magnitude of results were similar when excluding that year from the analyses (Appendix S1, Fig. S9).

### 1.3 Criteria to assess breeding status on transect

Reproductive status of individuals detected on transects was determined from their behaviour. For plovers and sandpipers, distraction displays (broken wing, rodent run) and insistent calls indicated nesting individuals (Lamarre *et al*. 2017), with status confirmed by approaching the bird. Individuals that did not react or were foraging, flying by, or resting, were considered non-breeders. Jaegers observed as one or two individuals along a transect were classified as nesting pair, as they typically leave their territory if breeding fails, while non-incubating groups of three or more were considered non-breeders (Andersson 1976). Snow geese were considered breeders when near a nest and non-breeders when in large groups or without a nest in sight. All detected Lapland longspurs, horned larks, and snow buntings were assumed to be nesting, as they exhibited singing, display flights, or alarm calls when approached (Drury 1961, Hunt et al. 1995).

### 1.4 Home range analysis

Between 2008 and 2023, we captured 165 foxes within the study area and equipped them with ARGOS or GPS collars (Clermont *et al*. 2021; Lai *et al*. 2015). We measured their annual home range (95% isopleth) using autocorrelated kernel density estimation implemented in the ctmm R package (Calabrese et al., 2016) as described in (Dulude-de Broin *et al*. 2023). The distribution of fox home range sizes was clearly bimodal with a large part of the variation explained by the presence of the goose colony (marginal R^2^=0.20, conditional R^2^= 0.46, n=248, from a linear mixed-effects model with home range size as the response variable, den location (inside vs outside of the colony) as fixed effect, and year as random effect). We therefore classified transects and lakes into two categories of predation risk to reflect this distribution. Using a Gaussian Mixture Model fitted with the mclust R package, we identified two distinct groups separating transects and lakes located within smaller home ranges (11 to 16.35 km²) corresponding to areas with high fox density, and larger home ranges (16.35 to 31 km²) corresponding to areas with lower fox density. In the smaller home range group, 85% of transects and lakes were within home ranges smaller than 15 km², while in the larger home range group, 85% were in home range larger than 18 km². To further validate our classification, we compared the occurrence of foraging foxes on transects in large and small home range, corresponding to low and high fox densities respectively. Occurrence of foraging foxes on transect were 4.2 times higher in areas of high fox densities compared to areas with low fox densities (Figure S1).

**Figure S1.**
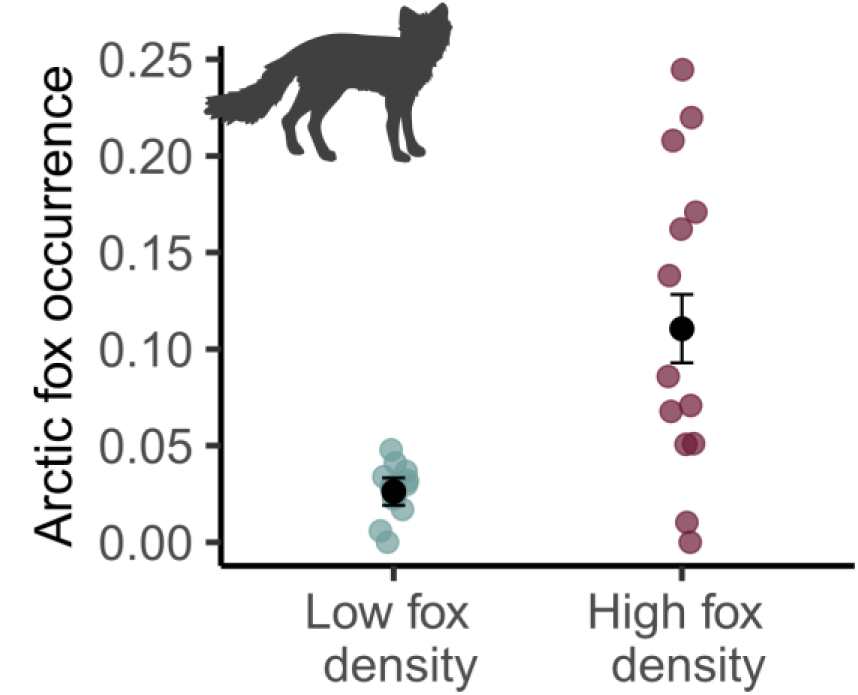
Arctic fox occurrence on transects in areas classified as low and high fox density on Bylot Island, Nunavut (2010–2024). Black dots and error bars represent average occurrence with 95% confidence intervals based on a binomial model with year and transect ID as random intercepts. Colored dots represent the proportion of fox sightings based on all transects for each fox density category in each year (n=30).

### 1.5 Bootstrap confidence intervals for snowmelt threshold

Snowmelt thresholds were defined for each species as the latest snowmelt date across all transects or lakes with a nesting individual. To obtain a measure of variability for this value, we resampled the entire dataset 500 times with replacement, calculated the threshold for each resample, and used the median as the final threshold with 95% quantiles as confidence intervals. A species with only one nesting observation near the threshold and no others close in time would yield a wide confidence interval. In contrast, if multiple transects or lakes with nesting individuals shared the same latest snowmelt date, the threshold would be more robust, resulting in a narrower confidence interval.

### 1.6 Number of surveys with sighting per species

Among nesting species, we recorded 3579 transect-years with Lapland longspurs, 1585 with snow geese, 1096 with American golden plovers, 629 with glaucous gulls, 530 with long-tailed jaegers, 468 with horned larks, 303 with Baird’s sandpipers, 296 with white-rumped sandpipers, 244 with snow buntings. Around lakes, we recorded 124 lake-years with cackling goose nests, 54 with red-throated loon and 37 with glaucous gulls. We recorded the presence of foraging arctic fox on 185 transects.

## 2. Enhanced snowmelt map from the integration of MODIS and Sentinel-2

To combine MODIS and Sentinel-2 derived products, we first extracted MODIS and Sentinel-2 data (section 2.1) and assessed the consistency of snowmelt date estimates from each method and satellite platform (section 2.2). Next, we calculated the difference in snowmelt dates between each MODIS pixel and all corresponding Sentinel-2 for the years when both datasets were available (n=5; 2018-2023) to generate a correction raster (section 2.3). We then added the median Sentinel-2-MODIS difference at each pixel to MODIS derived snowmelt rasters to generate a final 20 m resolution snowmelt map of all years. Finally, we validated our enhanced MODIS product against the original Sentinel-2 snowmelt maps for years when Sentinel-2 products were available (section 2.4).

### 2.1. Sentinel-2 and MODIS derived snow products

We obtained Sentinel-2 derived snowmelt dates (years 2017-2023) from the Sentinel-2 Level 3A snow product developed by THEIA (https://www.theia-land.fr/ces-cryosphere/neige/). This product uses all available Sentinel-2 images to track snow cover at a 20 m resolution and fills gaps caused by clouds or missing images using temporal interpolation (Barrou Dumont *et al*. 2025). Each image is analyzed to classify every pixel as either snow-covered or snow-free based on the Normalized Difference Snow Index (NDSI). The interpolated time series of these classifications is then used to estimate the snowmelt date, defined as the last day of the longest uninterrupted snow-covered period within the hydrological year. Compared to in situ measurements in the Alps and the Pyrenees, the method yielded a mean absolute error of 8 days, an root mean square error (RMSE) of 23 days, and a bias of -1 day (Barrou Dumont *et al*. 2025). Given the shorter Sentinel-2 revisit time in the high arctic (1-2 days) compared to these mid-latitude regions (5 days), errors are expected to be lower within our study area.

We obtained MODIS derived snowmelt dates (years 2010-2024) using MOD10A1 snow cover product Version 61 (Hall & Riggs 2021). This provides snow cover at a 500 m spatial resolution. To determine snowmelt dates for each year, we analyzed the snow cover time series for each pixel from March 1 to August 31. A pixel was considered snow-free for values <50, and the snowmelt date was defined as the first snow-free day followed by at least five consecutive snow-free days. To account for brief spring snowfall events that melt shortly after precipitation, we allowed a single day within this five-day period to exceed 50. Gaps in the time series, caused by clouds or missing imagery, were filled using linear interpolation. Detailed information on the MOD10A1 Snow Cover Product, Version 61, is provided in Hall et al. (2019).

### 2.2 Consistency between Sentinel-2 and MODIS derived snowmelt dates

We evaluated the relationship between snowmelt dates derived from MODIS (lower spatial, higher temporal resolution) and Sentinel-2 imagery (higher spatial, lower temporal resolution). Sentinel-2 maps were first aggregated to MODIS resolution (500 × 500 m) by averaging all corresponding high-resolution (20 × 20 m) Sentinel-2 pixels within each coarser pixel (Figure S2). This created a new 500 m x 500 m raster where each pixel reflects the aggregated value of the finer-scale data.

**Figure S2.**
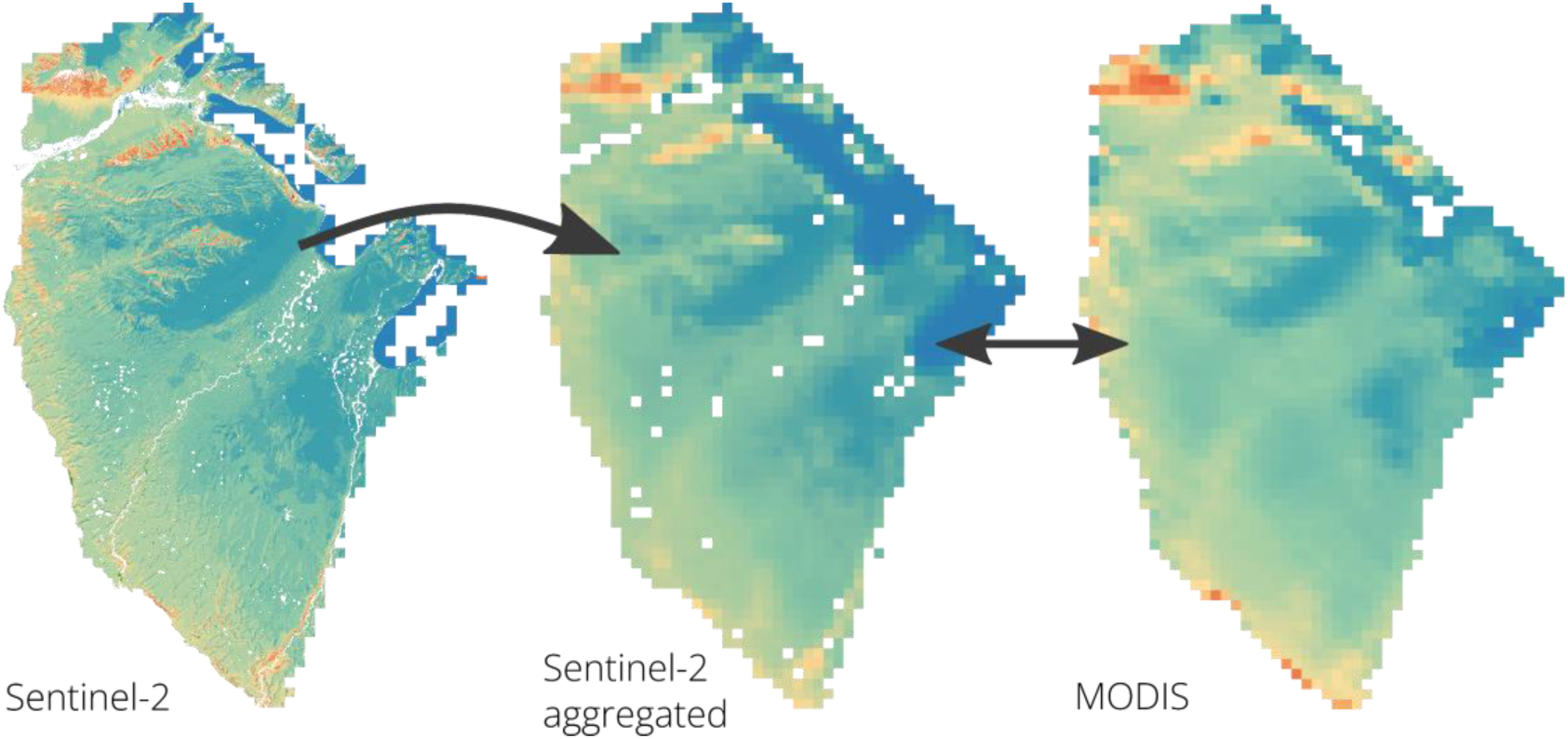
Comparison of Sentinel-2 and MODIS snow products for 2020. The Sentinel-2 map was aggregated to match MODIS spatial resolution (500 × 500 m) by averaging all corresponding high-resolution (20 × 20 m) Sentinel-2 pixels within each coarser pixel.

We then performed a pixel-by-pixel comparison of snowmelt dates at the 500 x 500 m resolution across five years (2018–2023) where both datasets were available. Snowmelt dates derived from Sentinel-2 and MODIS were highly related for all years with overlapping data (Figure S3; all slopes = 1.0, R² > 0.89, n=1618). The strong agreement between the two datasets, derived from different satellite platforms and analytical methods, suggests that snowmelt date estimates are consistent and support the combination of these snow products.

**Figure S3.**
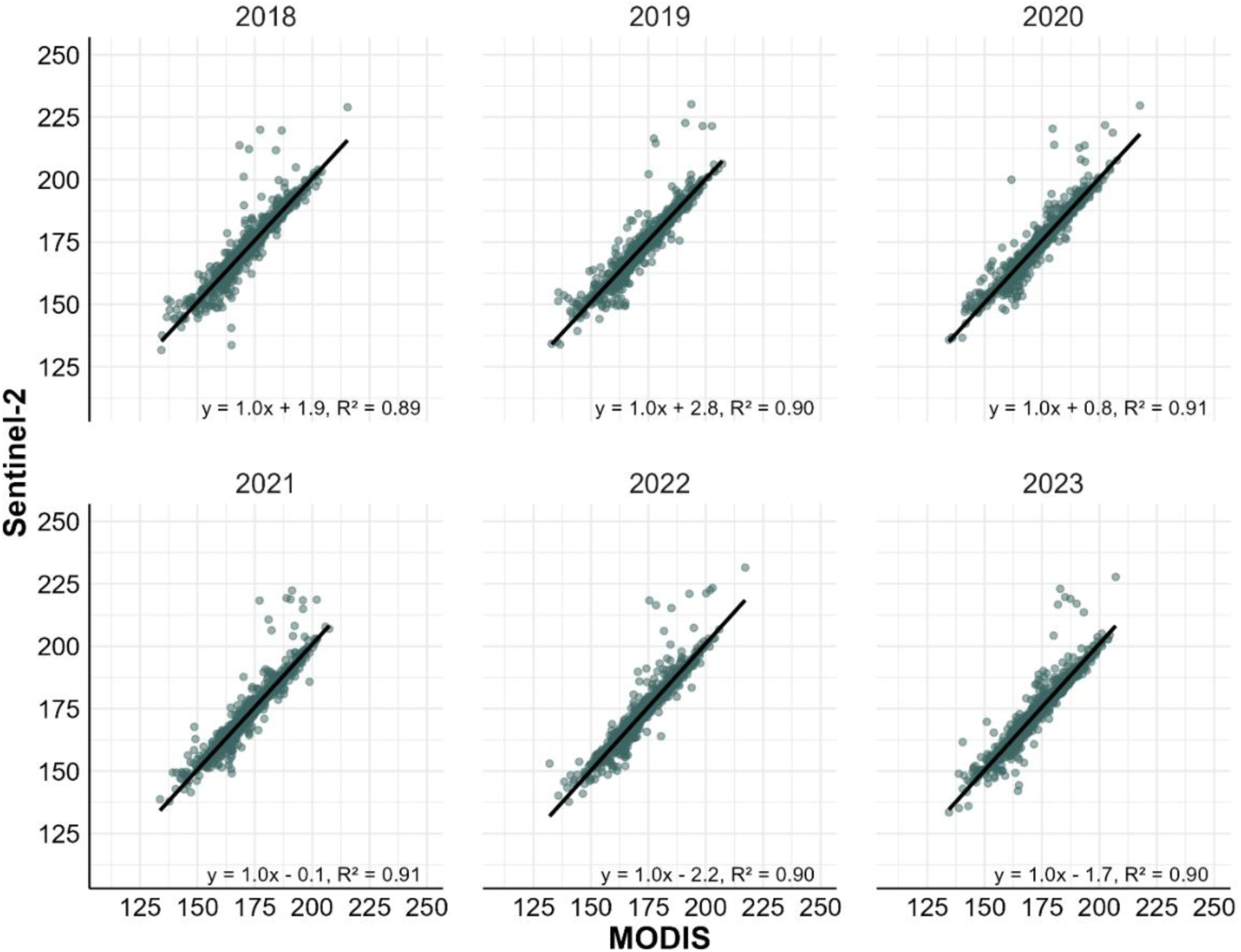
Relationship between snowmelt dates derived from Sentinel-2 and MODIS snow products. Snowmelt dates were compared on a pixel-by-pixel basis after aggregation of Sentinel-2 snowmelt data to match MODIS resolution. Linear model sample size: n=1618.

### 2.3 Creation of an adjustment raster to upscale MODIS product

To enhance the spatial resolution of MODIS snow products, we generated a correction raster from the difference between MODIS and Sentinel-2 derived snow rasters (Figure S4). For each 20 x 20 m pixel in the study area, we subtracted the value of the MODIS derived snowmelt date to the Sentinel-2 derived snowmelt date. As an illustration, if a large MODIS pixel indicated snowmelt on day 180, corresponding higher-resolution Sentinel-2 pixels within that area might show snowmelt occurring slightly earlier or later such as day 178, 184, or 174, resulting in difference values of –2, +4, and –6 days, respectively. These values were calculated for all 20 x 20 m pixels every year to generate the correction raster for all years when both data sources were available. Most annual differences in snowmelt dates at the 20 x 20 m resolution ranged between -20 and +20 days (figure S5), and interannual repeatability in adjustment values was high (Figure S6). We took the median value across years for each pixel to build the final adjustment raster. Median adjustment values ranged between -10 and + 10 days (Figure S5).

**Figure S4.**
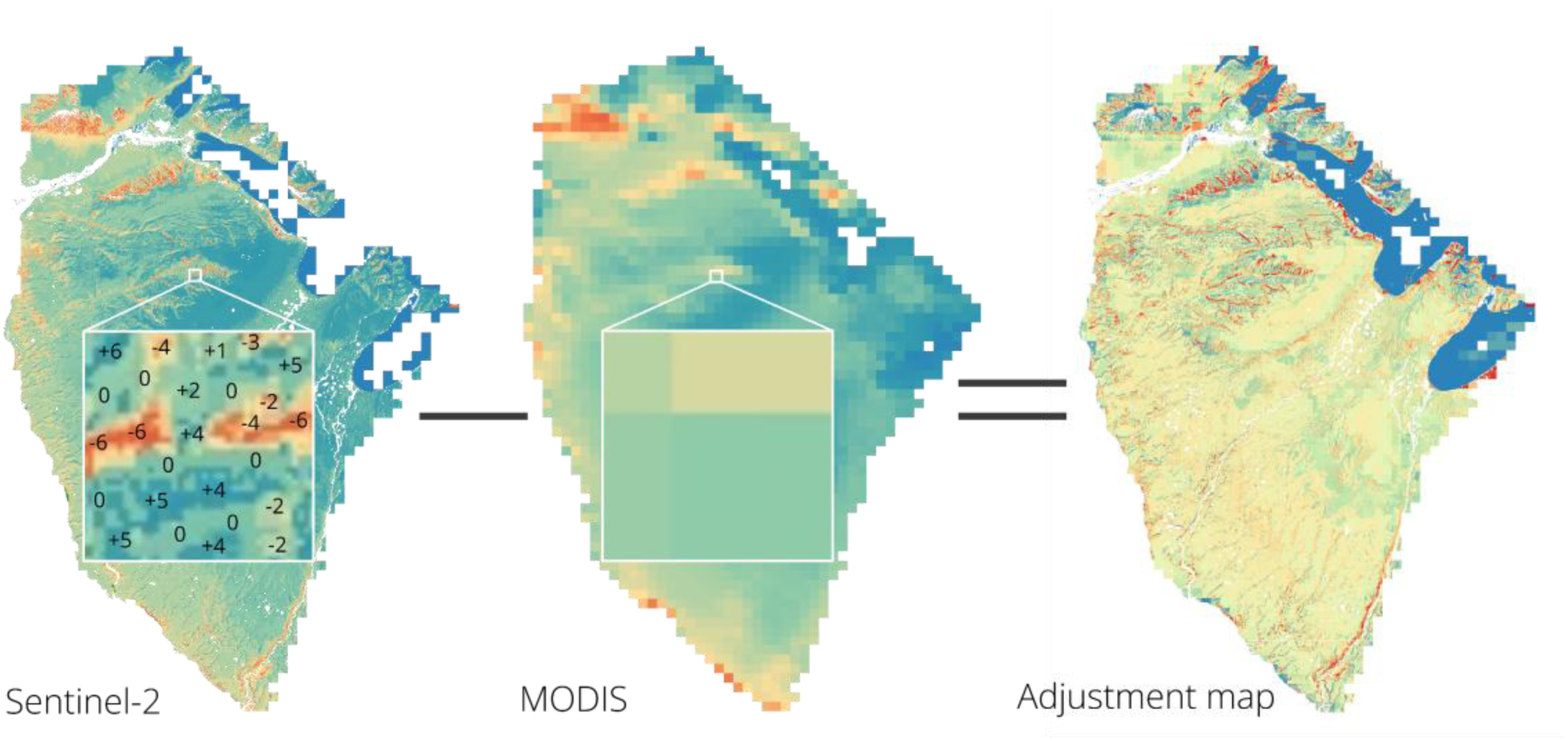
Difference between MODIS snowmelt dates and Sentinel-2 snowmelt dates to create the adjustment raster. The inset illustrates how high-resolution Sentinel-2 pixels differ from the corresponding coarser MODIS pixels. These differences constitute the values of the adjustment map.

**Figure S5.**
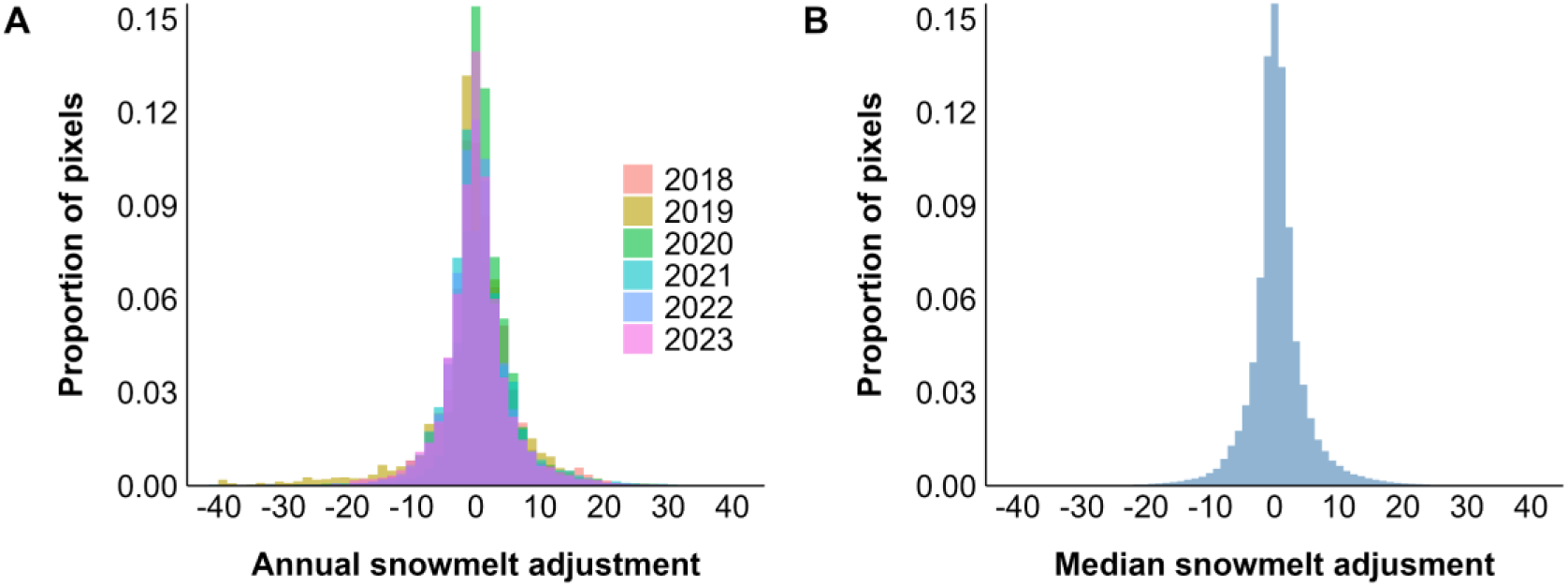
Distribution of snowmelt date adjustment values used to increase the spatial resolution of MODIS-derived snowmelt dates. Adjustment values correspond to the number of days difference between snowmelt estimates from each small Sentinel-2 pixel and coarser overlapping MODIS pixel. Annual adjustment values (A) were calculated for every 20 x 20 m pixel of the study area across all years where both data sources were available. The median difference per pixel (B) was used to create the final adjustment raster.

**Figure S6.**
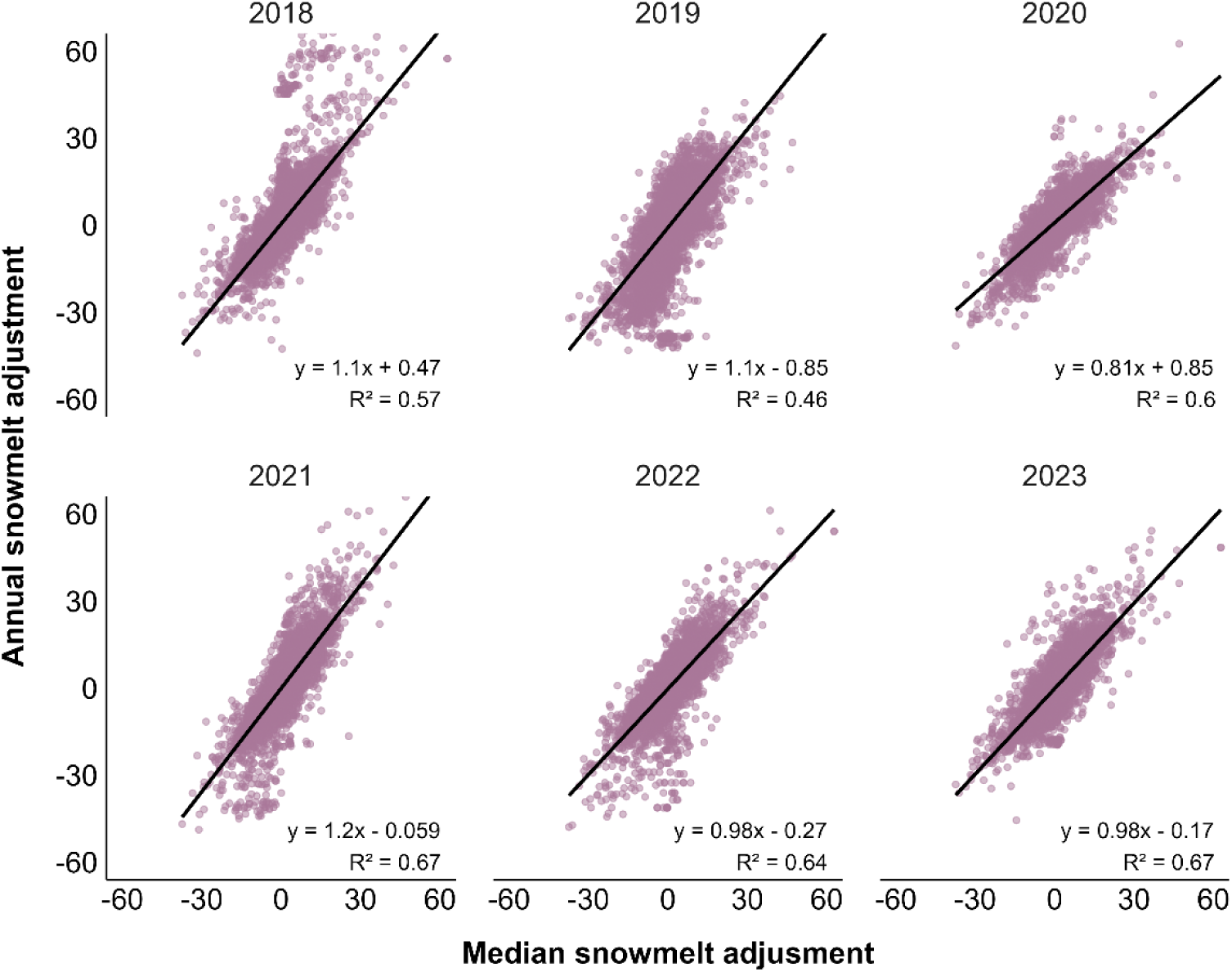
Repeatability in snowmelt adjustment values at the pixel level between Sentinel-2 and MODIS derived snow products. Adjustment values correspond to the number of days difference between snowmelt estimates from each small Sentinel-2 pixel and coarser overlapping MODIS pixel. Individual graphs relate the annual adjustment value to the median adjustment value across all years for the same pixel.

### 2.4 Final snowmelt map and validation against Sentinel-2

We produced the final snowmelt date maps by adding the adjustment layer to the MODIS-derived product (Figure S7), resulting in 20 m resolution snowmelt maps for all years from 2010 to 2024. To evaluate the accuracy of this hybrid product, we compared it against Sentinel-2 snowmelt maps for the years when both datasets were available (2018–2023). For both layers, we extracted the median snowmelt date within a 150 m buffer on each side of our transects (same as the main analysis). We then assessed the relationship between the enhanced MODIS and Sentinel-2 maps using a linear model (Figure S8-S9). The two products were highly related (Estimate [95% CI] = 0.99 [0.98, 1.00], n = 3298, R² = 0.93). The strong agreement is notable given that the adjustment map is the same for all years as it was obtained from median adjustment values. This further suggests that fine-scale spatial patterns in snowmelt timing were consistent across years (as previously shown in section 2.3, Figure S6), allowing accurate adjustment of MODIS data. The approach allowed us to obtain high-resolution and reliable snowmelt date maps for the entire study area across all years.

**Figure S7.**
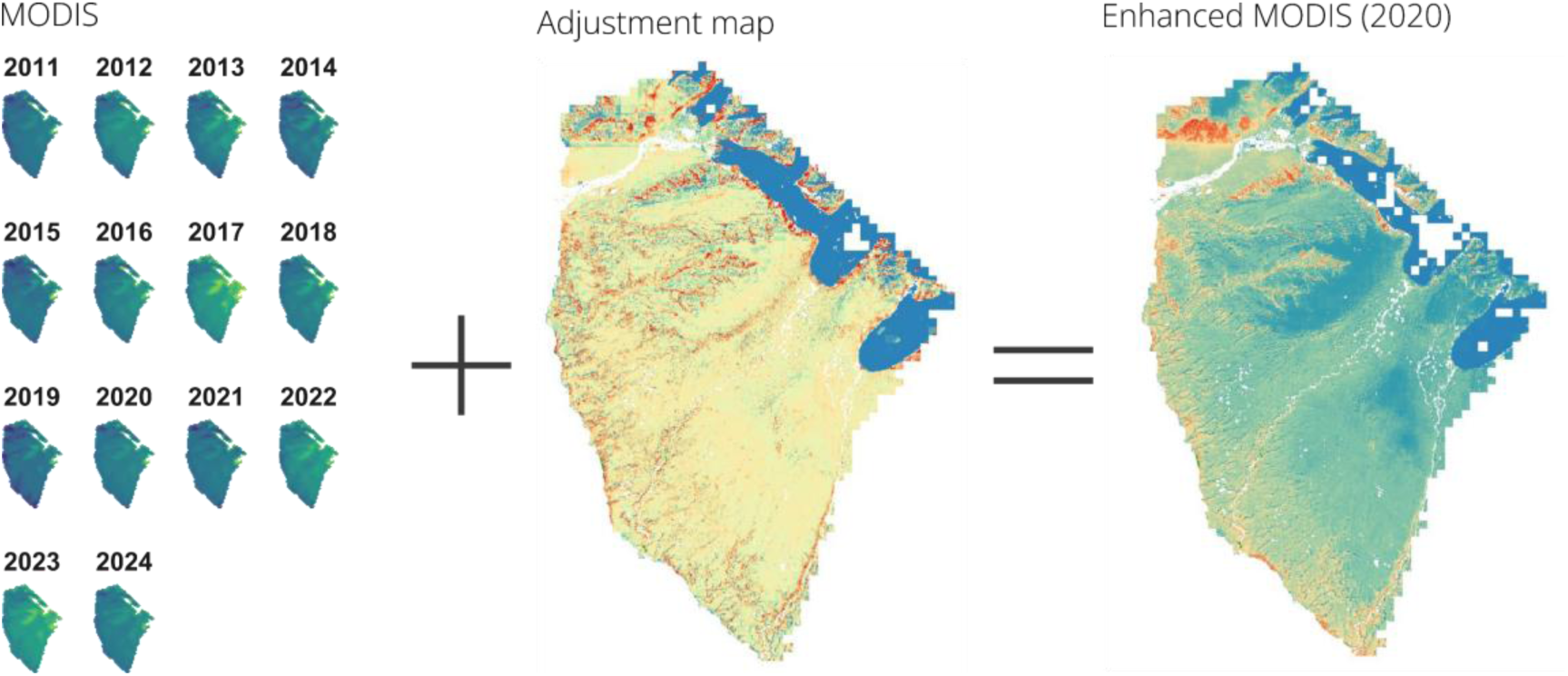
Method used to generate the final snow product used in the study. For each year, we adjusted the MODIS-derived snowmelt map using the adjustment map derived from Sentinel-2 data to obtain a product with a 20-m resolution.

**Figure S8.**
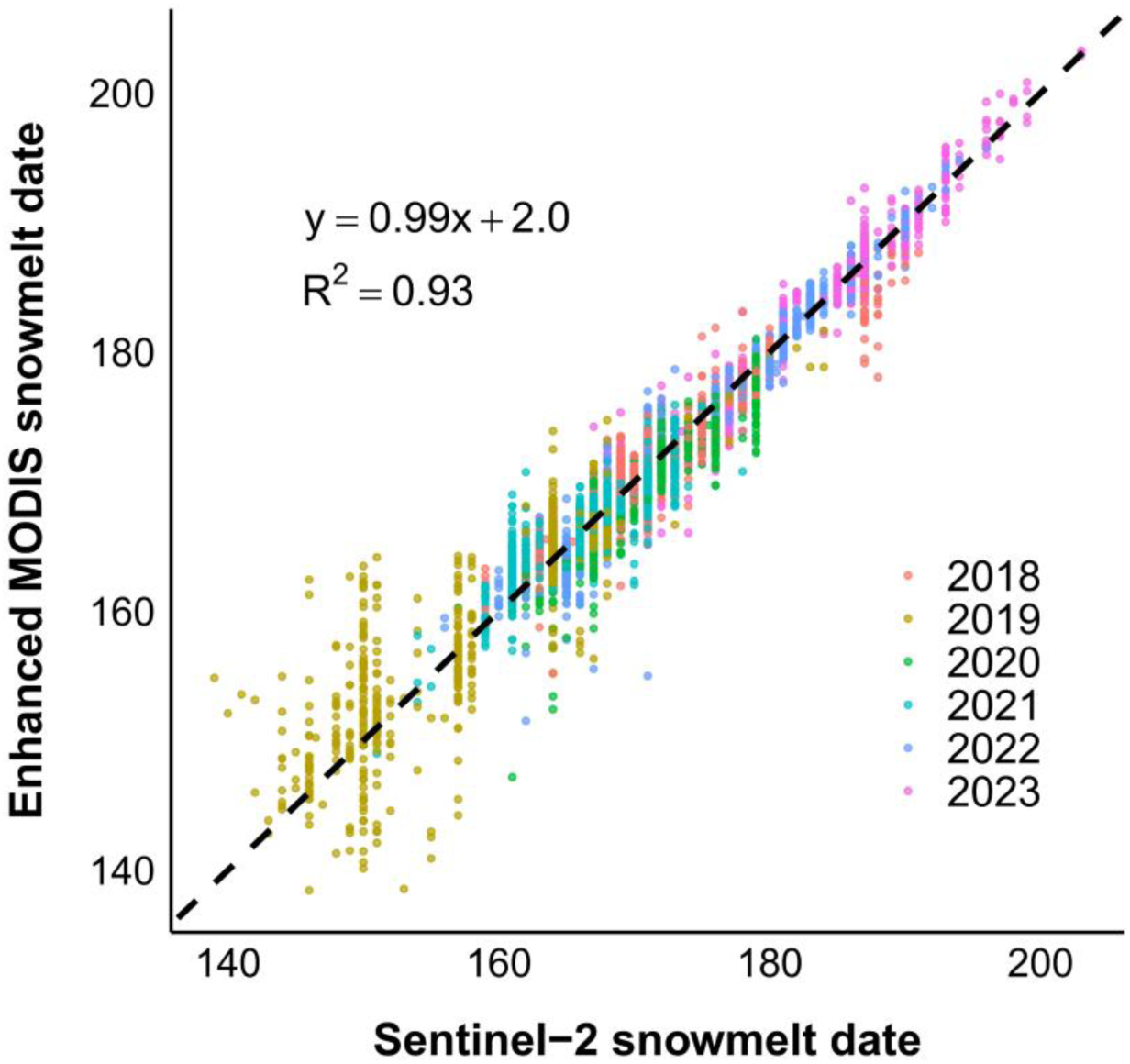
Validation of the snowmelt date derived from the enhanced MODIS snowmelt map against Sentinel-2 data for years with available Sentinel-2 coverage. Each point represents the median snowmelt date for each transect surveyed in the study area every year. The dashed line shows the 1:1 reference line. Dates are day of the year. Linear model sample size: n=3956.

**Figure S9.**
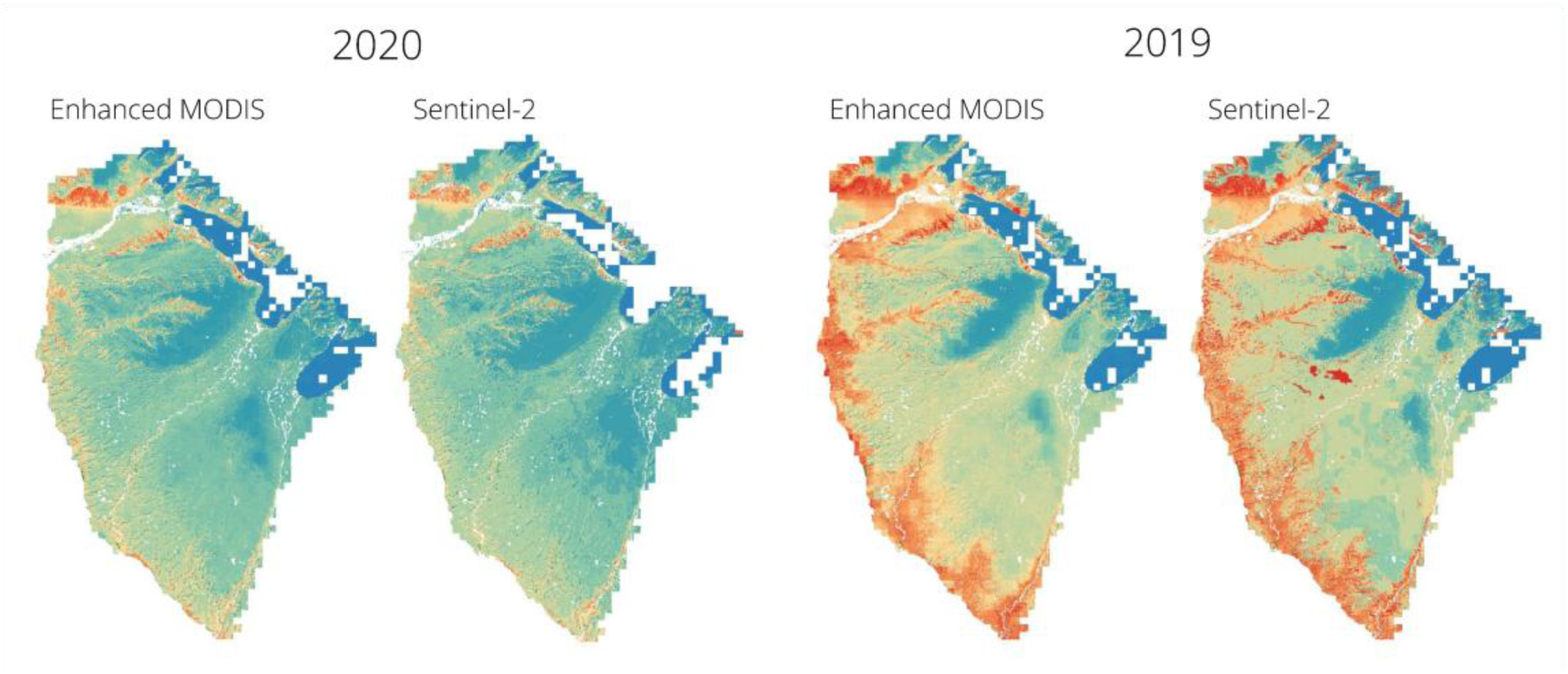
Visual comparison of enhanced MODIS and Sentinel-2–derived snowmelt products for two years with contrasting snowmelt timing, illustrating their consistency.

## 3. Relationship between the number of snow-free days and snowmelt date

We assessed the relationship between the number of snow-free days and snowmelt date using Sentinel-2 derived products from THEIA between 2018 and 2023. We extracted the median snow-free period (number of days between snowmelt and snow onset) and the median snowmelt date within a 150 m buffer (same as the main analysis) on each side of our 500 m transects. We then used a linear model to evaluate the relationship between the length of the snow-free period and snowmelt date. Snowmelt date was a strong predictor of the number of days without snow (Estimate [95%CI] = –1.05 [–1.08, –1.03], n = 3298, R² = 0.65, Figure S10), indicating that in addition to being the primary driver of primary production, snowmelt date is a good indicator of the duration of the snow-free window.

**Figure S10.**
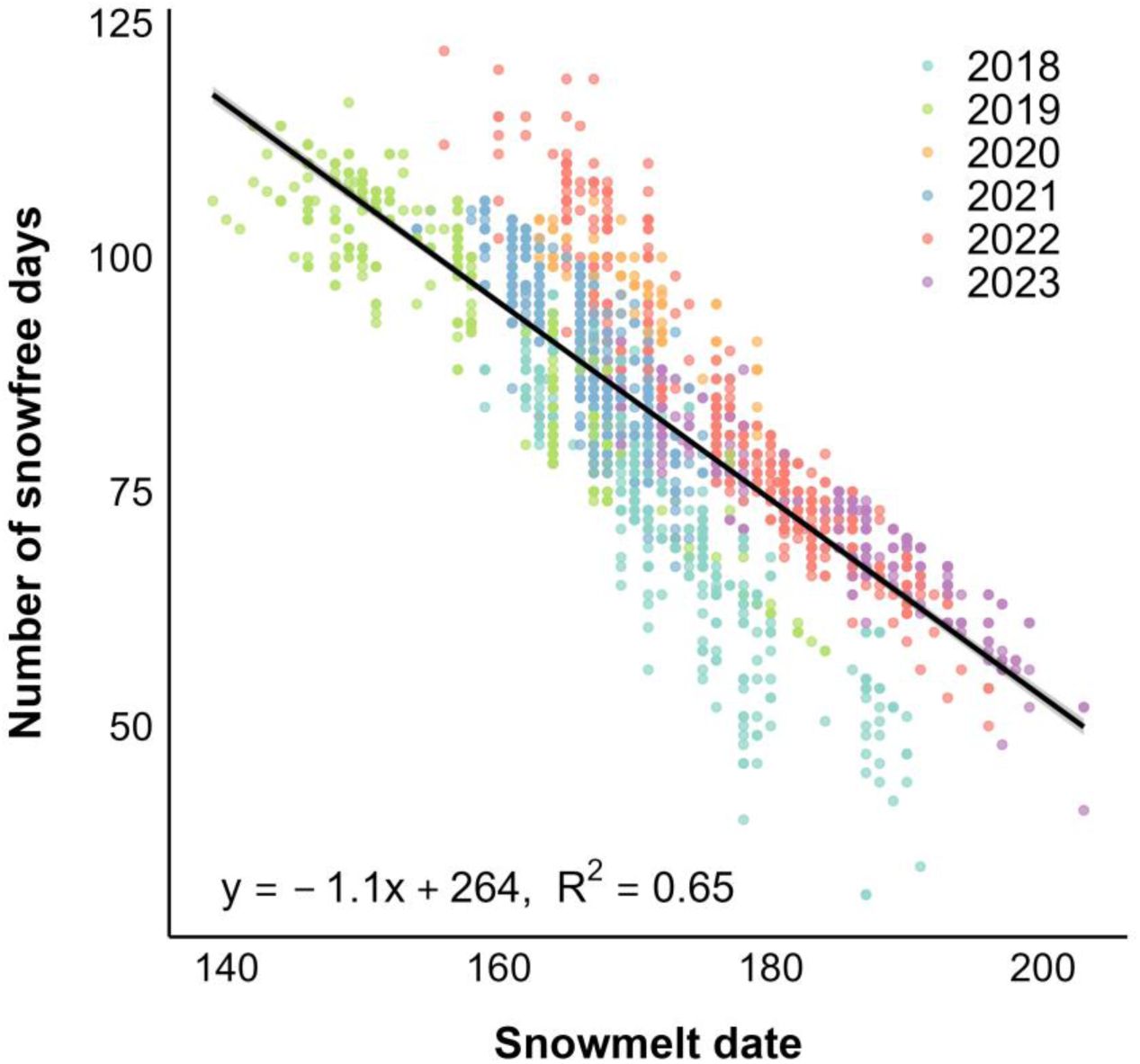
Relationship between snowmelt date and the number of snow-free days around individual transects every year, based on Sentinel-2 derived snow products from THEIA (2018–2023). Linear model sample size: n=3298.

## 4. Snowmelt threshold values by species

**Table S1.** Snowmelt threshold values for nesting birds on Bylot Island, Nunavut (2010–2024). Snowmelt threshold corresponds to the last date at which a breeding individual was sighted. Confidence intervals were obtained through non-parametric bootstrap.

| Species | Day of year | 95% CI | Calendar date<br>(non-leap year) |
| --- | --- | --- | --- |
| Greater snow goose | 184 | 183–184 | July 3 |
| Cackling goose | 179 | 178–179 | June 28 |
| Glaucous gull | 180 | 178–181 | June 29 |
| Long-tailed jaeger | 180 | 179–181 | June 29 |
| Red-throated loon | 189 | 187–190 | July 8 |
| American golden plover | 188 | 187–189 | July 7 |
| White-rumped sandpiper | 191 | 186–193 | July 10 |
| Baird’s sandpiper | 192 | 191–193 | July 11 |
| Lapland longspur | 197 | 194–198 | July 16 |
| Horned lark | 199 | 196–199 | July 18 |
| Snow bunting | 201 | 197–201 | July 20 |

## 5. Model selection results for all species

**Table S2.**
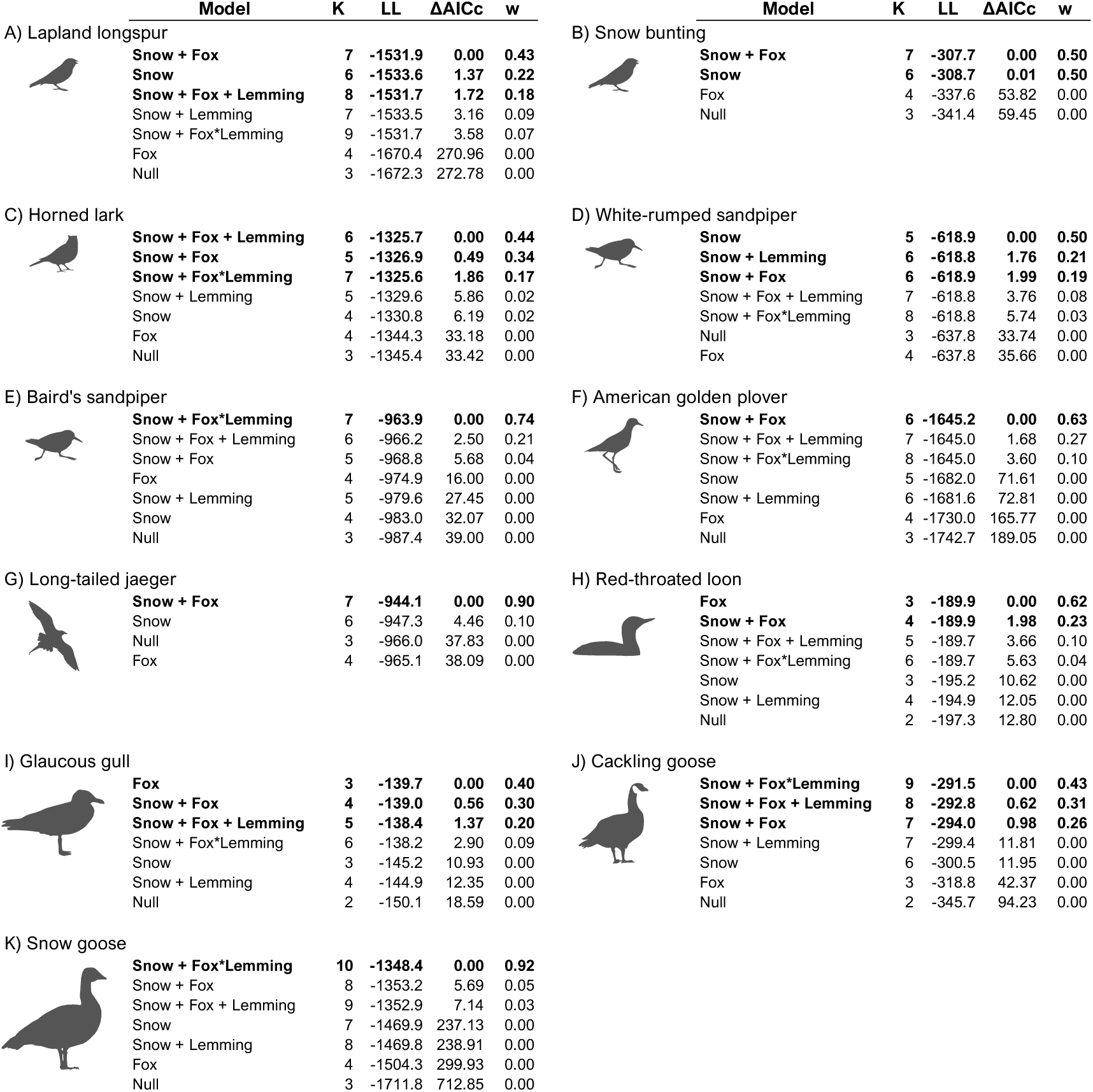
Model selection results for breeding species occurrence in relation to snowmelt date, fox density, and lemming density. All models are generalized linear mixed models with a binomial distribution and include year as a random intercept. Models for tundra-nesting birds (species A–G and K) are based on presence/absence data from 500 m transects and include transect ID as a random intercept. Models for lake-associated birds (species H–I) are based on presence/absence data around lakes. Models are ranked by Akaike Information Criterion (AICc). The table shows the number of parameters (K), log-likelihood (LL), delta AICc (ΔAICc), and AICc weight (w). Models in bold are considered competitive (delta AICc < 2).

| | Model | K | LL | $\Delta AICc$ | w |
| --- | --- | --- | --- | --- | --- |
| A) Lapland longspur |  |  |  |  |  |
|  | **Snow + Fox** | 7 | -1531.9 | 0.00 | 0.43 |
|  | **Snow** | 6 | -1533.6 | 1.37 | 0.22 |
|  | **Snow + Fox + Lemming** | 8 | -1531.7 | 1.72 | 0.18 |
|  | Snow + Lemming | 7 | -1533.5 | 3.16 | 0.09 |
|  | Snow + Fox*Lemming | 9 | -1531.7 | 3.58 | 0.07 |
|  | Fox | 4 | -1670.4 | 270.96 | 0.00 |
|  | Null | 3 | -1672.3 | 272.78 | 0.00 |
| B) Snow bunting |  |  |  |  |  |
|  | **Snow + Fox** | 7 | -307.7 | 0.00 | 0.50 |
|  | **Snow** | 6 | -308.7 | 0.01 | 0.50 |
|  | Fox | 4 | -337.6 | 53.82 | 0.00 |
|  | Null | 3 | -341.4 | 59.45 | 0.00 |
| C) Horned lark |  |  |  |  |  |
|  | **Snow + Fox + Lemming** | 6 | -1325.7 | 0.00 | 0.44 |
|  | **Snow + Fox** | 5 | -1326.9 | 0.49 | 0.34 |
|  | **Snow + Fox\*Lemming** | 7 | -1325.6 | 1.86 | 0.17 |
|  | Snow + Lemming | 5 | -1329.6 | 5.86 | 0.02 |
|  | Snow | 4 | -1330.8 | 6.19 | 0.02 |
|  | Fox | 4 | -1344.3 | 33.18 | 0.00 |
|  | Null | 3 | -1345.4 | 33.42 | 0.00 |
| D) White-rumped sandpiper |  |  |  |  |  |
|  | **Snow** | 5 | -618.9 | 0.00 | 0.50 |
|  | **Snow + Lemming** | 6 | -618.8 | 1.76 | 0.21 |
|  | **Snow + Fox** | 6 | -618.9 | 1.99 | 0.19 |
|  | Snow + Fox + Lemming | 7 | -618.8 | 3.76 | 0.08 |
|  | Snow + Fox*Lemming | 8 | -618.8 | 5.74 | 0.03 |
|  | Null | 3 | -637.8 | 33.74 | 0.00 |
|  | Fox | 4 | -637.8 | 35.66 | 0.00 |
| E) Baird's sandpiper |  |  |  |  |  |
|  | **Snow + Fox\*Lemming** | 7 | -963.9 | 0.00 | 0.74 |
|  | Snow + Fox + Lemming | 6 | -966.2 | 2.50 | 0.21 |
|  | Snow + Fox | 5 | -968.8 | 5.68 | 0.04 |
|  | Fox | 4 | -974.9 | 16.00 | 0.00 |
|  | Snow + Lemming | 5 | -979.6 | 27.45 | 0.00 |
|  | Snow | 4 | -983.0 | 32.07 | 0.00 |
|  | Null | 3 | -987.4 | 39.00 | 0.00 |
| F) American golden plover |  |  |  |  |  |
|  | **Snow + Fox** | 6 | -1645.2 | 0.00 | 0.63 |
|  | Snow + Fox + Lemming | 7 | -1645.0 | 1.68 | 0.27 |
|  | Snow + Fox*Lemming | 8 | -1645.0 | 3.60 | 0.10 |
|  | Snow | 5 | -1682.0 | 71.61 | 0.00 |
|  | Snow + Lemming | 6 | -1681.6 | 72.81 | 0.00 |
|  | Fox | 4 | -1730.0 | 165.77 | 0.00 |
|  | Null | 3 | -1742.7 | 189.05 | 0.00 |
| G) Long-tailed jaeger |  |  |  |  |  |
|  | **Snow + Fox** | 7 | -944.1 | 0.00 | 0.90 |
|  | Snow | 6 | -947.3 | 4.46 | 0.10 |
|  | Null | 3 | -966.0 | 37.83 | 0.00 |
|  | Fox | 4 | -965.1 | 38.09 | 0.00 |
| H) Red-throated loon |  |  |  |  |  |
|  | **Fox** | 3 | -189.9 | 0.00 | 0.62 |
|  | **Snow + Fox** | 4 | -189.9 | 1.98 | 0.23 |
|  | Snow + Fox + Lemming | 5 | -189.7 | 3.66 | 0.10 |
|  | Snow + Fox*Lemming | 6 | -189.7 | 5.63 | 0.04 |
|  | Snow | 3 | -195.2 | 10.62 | 0.00 |
|  | Snow + Lemming | 4 | -194.9 | 12.05 | 0.00 |
|  | Null | 2 | -197.3 | 12.80 | 0.00 |
| I) Glaucous gull |  |  |  |  |  |
|  | **Fox** | 3 | -139.7 | 0.00 | 0.40 |
|  | **Snow + Fox** | 4 | -139.0 | 0.56 | 0.30 |
|  | **Snow + Fox + Lemming** | 5 | -138.4 | 1.37 | 0.20 |
|  | Snow + Fox*Lemming | 6 | -138.2 | 2.90 | 0.09 |
|  | Snow | 3 | -145.2 | 10.93 | 0.00 |
|  | Snow + Lemming | 4 | -144.9 | 12.35 | 0.00 |
|  | Null | 2 | -150.1 | 18.59 | 0.00 |
| J) Cackling goose |  |  |  |  |  |
|  | **Snow + Fox\*Lemming** | 9 | -291.5 | 0.00 | 0.43 |
|  | **Snow + Fox + Lemming** | 8 | -292.8 | 0.62 | 0.31 |
|  | **Snow + Fox** | 7 | -294.0 | 0.98 | 0.26 |
|  | Snow + Lemming | 7 | -299.4 | 11.81 | 0.00 |
|  | Snow | 6 | -300.5 | 11.95 | 0.00 |
|  | Fox | 3 | -318.8 | 42.37 | 0.00 |
|  | Null | 2 | -345.7 | 94.23 | 0.00 |
| K) Snow goose |  |  |  |  |  |
|  | **Snow + Fox\*Lemming** | 10 | -1348.4 | 0.00 | 0.92 |
|  | Snow + Fox | 8 | -1353.2 | 5.69 | 0.05 |
|  | Snow + Fox + Lemming | 9 | -1352.9 | 7.14 | 0.03 |
|  | Snow | 7 | -1469.9 | 237.13 | 0.00 |
|  | Snow + Lemming | 8 | -1469.8 | 238.91 | 0.00 |
|  | Fox | 4 | -1504.3 | 299.93 | 0.00 |
|  | Null | 3 | -1711.8 | 712.85 | 0.00 |

## 6. Model results for all species

**Table S3.**
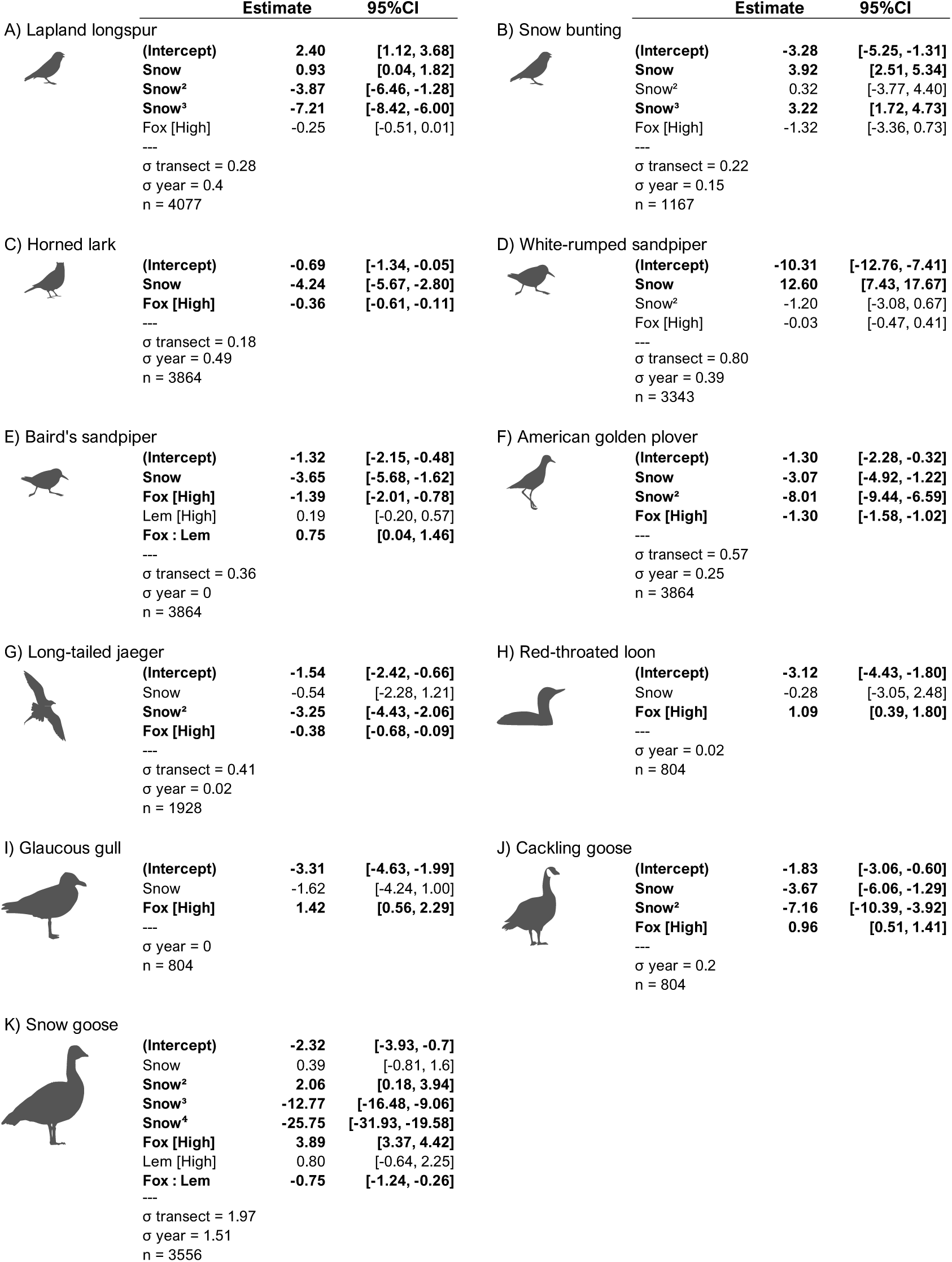
Parameter values for selected models relating breeding species occurrence to snowmelt date (Snow), fox density (Fox), and lemming density (Lem) in Table S2. Models for tundra-nesting birds (species A–G and K) and wetland-associated birds (species H–I) are based on transects and lakes survey respectively. To illustrate the effect, or lack of effect, of snowmelt date and fox density, we present results from the most parsimonious model including both variables. Values with 95% confidence intervals excluding zero are shown in bold.

## 7. Results obtained when excluding data from 2024

Results were unaffected when excluding data from 2024 (Table S4 and Figures S11, S12, S13), a year in which surveys were delayed due to helicopter availability. The only notable difference was the long-tailed jaeger for which we no longer had data in late snowmelt areas. This species nest only in years of high lemming density, which substantially reduces sample size, particularly in late snowmelt areas for which 2024 was our only year of high lemming abundance. In the range for which we had data with and without 2024, patterns for jaeger were unaffected by the removal of 2024 data. Results for all other species were highly consistent with and without 2024 data.

**Table S4.**
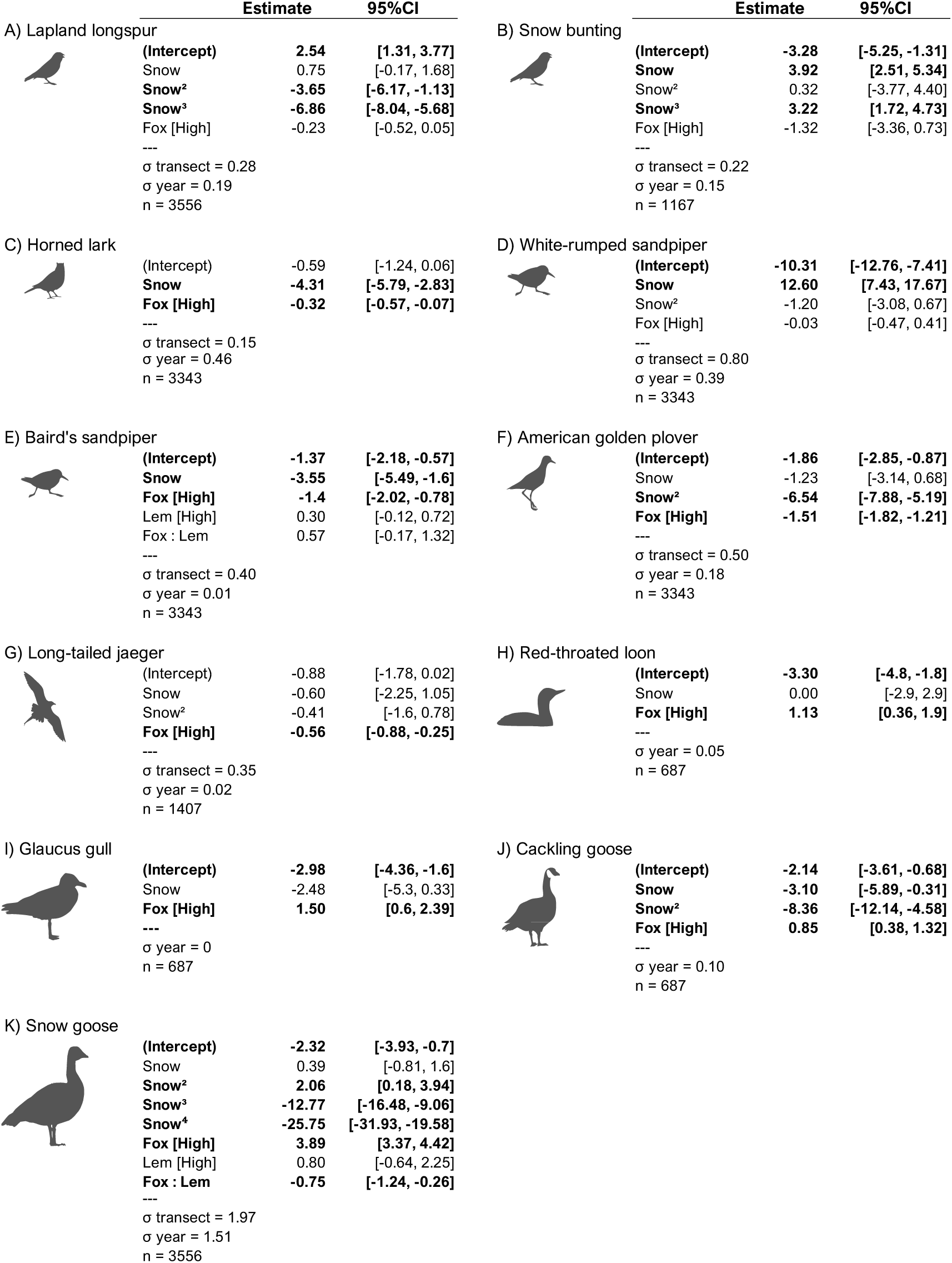
Model results for nesting species occurrence in relation to snowmelt date (Snow), fox density (Fox), and lemming density (Lem) when excluding data from 2024.

**Figure S11.**
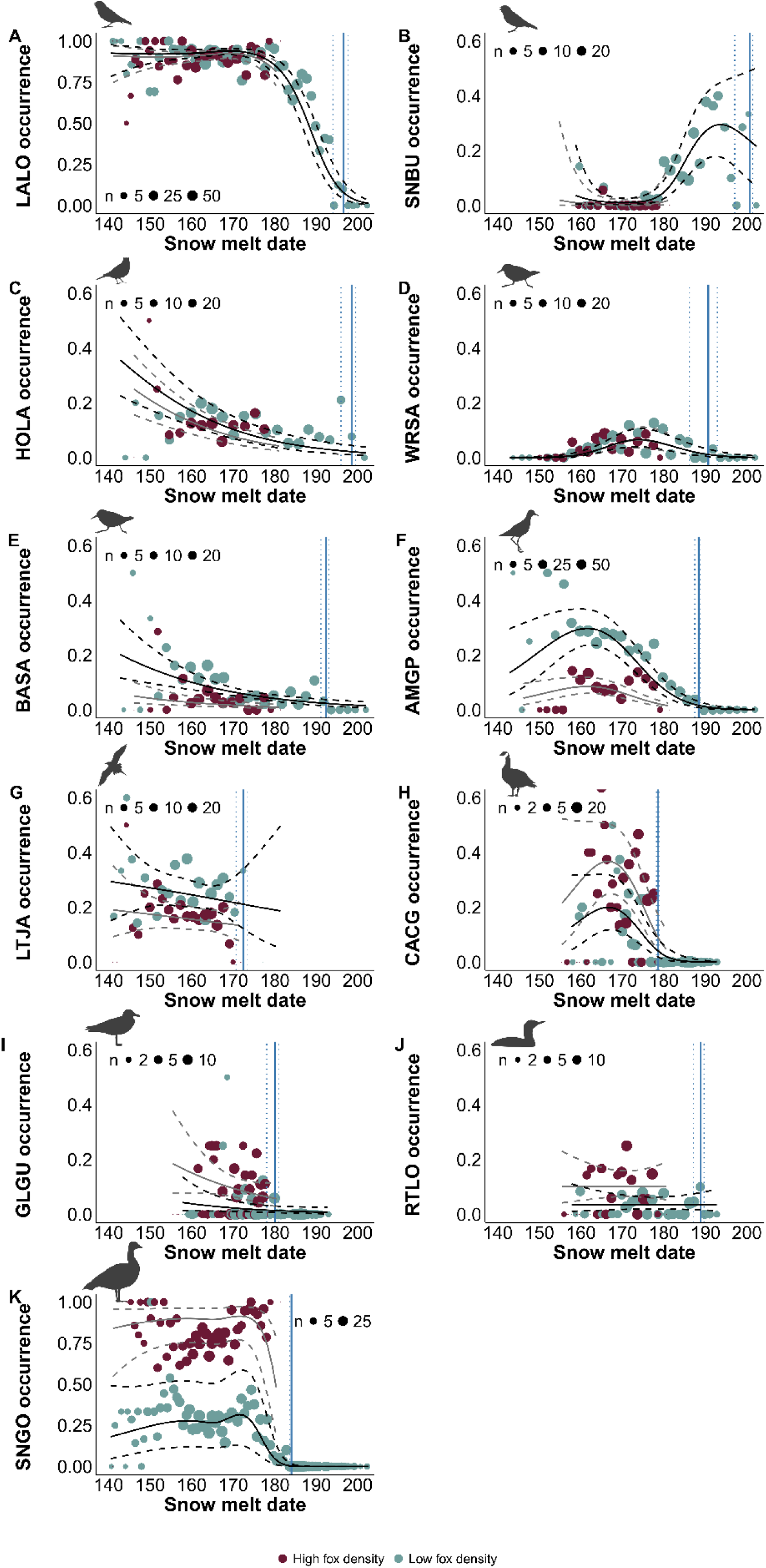
Influence of snowmelt date on occurrence probability of breeding species on Bylot Island, Nunavut when excluding data from 2024 (2010–2023).

**Figure S11.**
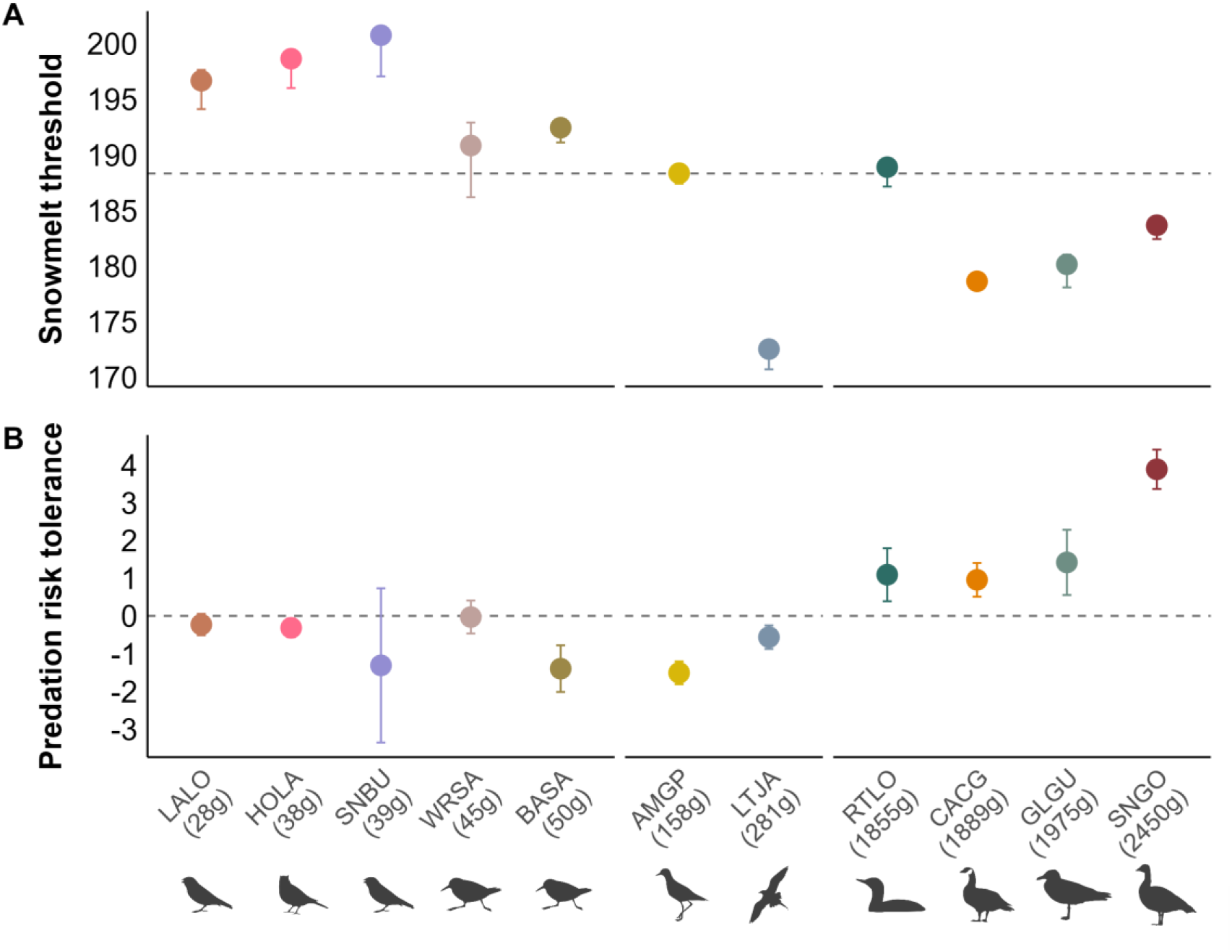
Snowmelt threshold dates for reproduction and predation risk tolerance of species ordered by body mass when excluding data from 2024. (A) Snowmelt threshold corresponds to the last date a breeding individual was observed. Dots show the threshold for each species with 95% confidence intervals from non-parametric bootstrap. (B) Predation risk tolerance is the effect of fox density on species occurrence probability (values in Table S4), with dots showing effect sizes and 95% confidence intervals. Negative predation risk tolerance indicates lower probability of occurrence in high fox density area.

## Notes

### Competing Interest Statement

The authors have declared no competing interest.

### Summary of Updates

Word count reduction and edits to figures

